# IRF8 configures enhancer landscape in postnatal microglia and directs microglia specific transcriptional programs

**DOI:** 10.1101/2023.06.25.546453

**Authors:** Keita Saeki, Richard Pan, Eunju Lee, Daisuke Kurotaki, Keiko Ozato

## Abstract

Microglia are innate immune cells in the brain. Transcription factor IRF8 is highly expressed in microglia. However, its role in postnatal microglia development is unknown. We demonstrate that IRF8 binds stepwise to enhancer regions of postnatal microglia along with Sall1 and PU.1, reaching a maximum after day 14. IRF8 binding correlated with a stepwise increase in chromatin accessibility, which preceded the initiation of microglia-specific transcriptome. Constitutive and postnatal *Irf8* deletion led to a loss of microglia identity and gain of disease-associated microglia-like genes. Combined analysis of scRNA-seq and scATAC-seq revealed a correlation between chromatin accessibility and transcriptome at a single-cell level. IRF8 was also required for microglia-specific DNA methylation patterns. Lastly, in the 5xFAD model, constitutive and postnatal *Irf8* deletion reduced the interaction of microglia with Aβ plaques and the size of plaques, lessening neuronal loss. Together, IRF8 sets the epigenetic landscape, which is required for postnatal microglia gene expression.

## Introduction

Microglia elicit innate immunity in the brain, providing antimicrobial protection^1, 2^. Microglia also promote brain development through synaptic pruning, supporting neuronal survival, and vascular formation^3, 4, 5, 6^. Microglia continuously surveil the entire brain and respond to changes in their environment^7, 8, 9^. Dysregulation of microglia is associated with neurodegenerative diseases such as Alzheimer’s disease (AD)^10, 11^.

Microglia arise from the yolk sac-derived progenitors and expand in the brain in SPI1 (PU.1) and IRF8-dependent manner^12^. Transcription factors, such as Smad3 and Mafb, contribute to microglial differentiation^13^. Evidence indicates that microglia mature in the postnatal stage since they do not elicit full gene expression until later^14, 15^. Postnatal microglia maturation may be part of global change in the brain to adapt to the extra-uterine environment and develop further into adulthood^6, 14, 15^. However, postnatal microglia development and mechanisms controlling the process have not been fully understood.

IRF8 is a DNA binding transcription factor that binds to GAAAG ISRE element (ISRE) and TTCC…G/CTTT PU.1/IRF composite motif (ETS/ISRE), the latter by interacting with PU.1^16^. Outside the brain, IRF8 is expressed in developing myeloid cells in bone marrow, Ly6C^+^ macrophages, and CD8^+^ or plasmacytoid dendritic cells^17, 18^. IRF8 is essential for innate resistance against various pathogens^19^. Constitutive *Irf8* knockout (IRF8KO) mice are susceptible to infection while having excess neutrophil-like cells^20^. In the brain, IRF8 is expressed only in microglia, except for a few macrophages^21^. Besides participating in early microglia development, IRF8 responds to cytokines and pain injury in adults^12, 22, 23^. However, the role of IRF8 in postnatal microglia maturation is largely unknown^13, 14^.

CUT&RUN analysis revealed that IRF8 binds to enhancer regions of postnatal microglia gradually from P9 to P14, reaching a maximum at adult. IRF8 binding was a prerequisite for setting chromatin accessibility, histone modifications, and DNA methylation patterns unique to microglia, which preceded microglia specific gene expression. Postnatal deletion of *Irf8* (IRF8cKO) demonstrated that continuous IRF8 expression after P14 is necessary for maintaining microglia gene expression. Lastly, in the 5xFAD mouse model of AD, constitutive and postnatal deletion of *Irf8* led to reduced AD pathology due to reduced interaction of microglia with amyloidβ (Aβ) plaques. Together, IRF8 defines the epigenetic landscape in postnatal microglia, thereby directing their transcriptome programs.

## Results

### IRF8 binds to the microglia genome during postnatal development in a stepwise fashion

The IRF8’s role in postnatal development and adult microglia remains unclear. To gain insight into this question, we sought to identify genome-wide IRF8 binding sites in postnatally developing microglia. To first verify IRF8 expression in postnatal microglia, we examined Irf8-GFP knock-in mice that express endogenous IRF8 fused to GFP^24^. FACS analyses of microglia from P9, P14, and P60 (adult) showed that IRF8 is expressed at similar levels at all times, higher than that in peritoneal macrophages (Fig. S1A, S1B).

CUT&RUN experiments were carried out for microglia isolated from Irf8-GFP mice using anti-GFP antibody. Peritoneal macrophages from the same mice were also examined as a reference. We identified 16,935 IRF8 peaks in adult microglia and 12,062 in peripheral peritoneal macrophages (Fig. 1A). About half of the IRF8 peaks were in the intergenic regions with >5 kb distal from the genes, while about 20% were at the promoter regions in both cell types (Fig. 1A). *De novo* motif analysis revealed enrichment of ETS/ISRE and ISRE motifs (Fig. 1B). DEseq2-based statistics revealed a differential IRF8 binding profile between microglia and peritoneal macrophages (Fig. 1C). Further, GREAT GO annotation indicated microgli-specific IRF8 peaks (n=9,679) to be related to cell/leukocyte activation and differentiation (Fig. 1D; upper panel). In contrast, macrophage-specific IRF8 peaks (n=2,251) were related to immune responses and inflammatory responses (Fig. 1D; lower panel). In addition, in microglia, IRF8 binding was found near microglia identity genes, such as *Sall1* and *Cx3cr1*^11, 25^, but not in peritoneal macrophages (Fig. 1G).

**Figure 1.**
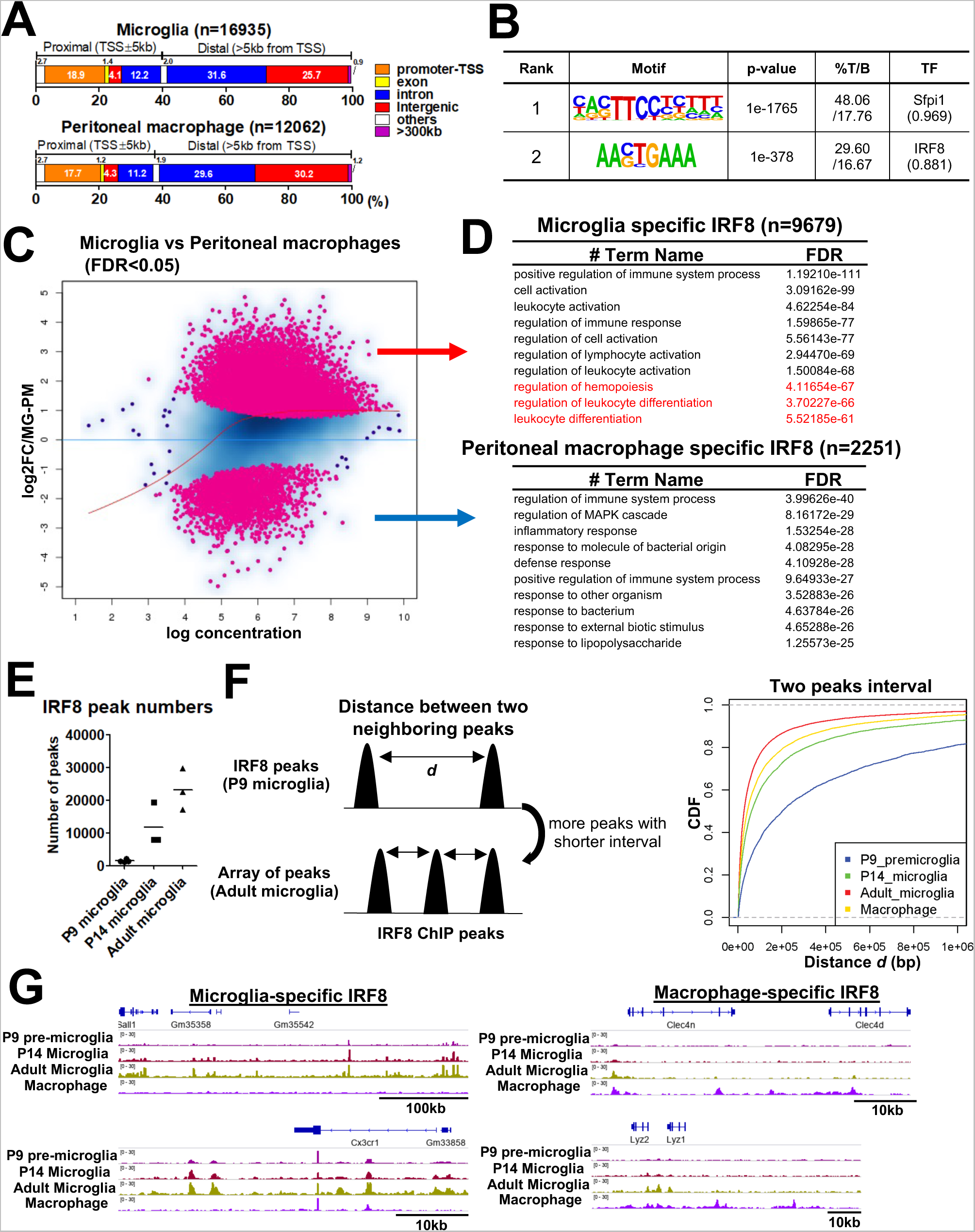
Microglia specific IRF8 occupancy increases postnatally. A. Genome wide distribution of IRF8 peaks in adult microglia (top) and peritoneal macrophages (bottom). The genomic position of each peak was determined with HOMER. B. HOMER *de novo* motif analysis for IRF8 binding sites in microglia. Motifs with a match score >0.85 are shown. A motif "Sfpi" is equivalent to the PU.1 motif. C. An M-A plot visualizing the difference between microglia IRF8 peak intensity and that of peritoneal macrophages (n=3). The differential peaks with FDR<0.05 are indicated in pink. D. Top 10 Gene ontology terms for microglia specific IRF8 peaks (top) and macrophage specific IRF8 peaks (bottom), analyzed with GREAT. Sequences covering 500kb distal from the TSS to 1kb from TES were set as an associating region. The terms of the GO biological process and the corresponding binomial FDR values were shown. E. The number of IRF8 peaks identified in P9, P14, and adult microglia. F. Schematic illustration of the distance between two neighboring IRF8 peaks; the shortening peak intervals represent an array-like appearance of peaks (left). The CDF plot shows that the frequency of short interval peaks increases along with the postnatal microglia development and more than peritoneal macrophages. (right; p=2.2×10^-16^, Kolmogorov-Smirnov two-sided Test). G. IGV examples of IRF8 peaks in developing and adult microglia and peritoneal macrophages at microglia specific genes (*Sall1* and *Cx3cr1*; left) or macrophage specific genes (*Clec4n*, *Lyz1*, and *Lyz2*; right).

IRF8 binding peaks on P9, P14, and adult microglia are shown in Fig. 1E. The IRF8 binding peaks on P9 microglia were very sparse. The binding increased at P14, although still less than on adult microglia, indicating a stepwise increase in IRF8 binding (Fig. 1E). Furthermore, cumulative distribution frequency graphs plotting the distance between two neighboring IRF8 binding sites (Fig. 1F, right) showed that the distance in the adult is the shortest, and the longest in P9 microglia, indicating that IRF8 binding in increases by forming arrays during postnatal development (Fig.1F left). IGV tracks corroborated the stepwise increase in IRF8 binding observed over the enhancer region of *Sall1* and *Cx3cr1* genes (Fig. 1G, left). In peritoneal macrophages, IRF8 was bound to regulatory regions of *Clec4n and Lyz1/2* which are active in these cell*s* (Fig. 1G, right). Thus, IRF8 binding to the microglial genome is developmentally controlled and reaches a maximum in adults when microglia specific genes become fully expressed^14, 15^. It is of note that despite a marked increase in binding activity, the amount of IRF8 was virtually the same during postnatal stages (Fig. S1B).

### IRF8 directs the expression of microglia identity genes and suppresses DAM genes

To study how IRF8 binding controls microglia’s transcriptional programs, we carried out bulk RNA-seq for microglia isolated from wild-type (WT) and constitutive IRF8KO mice. FACS analyses of brain cells revealed two CD45^+^ populations in the IRF8KO brain, one CD11b^high^CD45^+^ corresponding to WT microglia, and another CD11b^low^CD45^+^ subset, termed P3, which was absent in WT brain (Fig. S2A, S2B). The number of CD11b^high^CD45^+^ cells in IRF8KO brains was less than that of WT microglia. Cells termed P3, about 10 % of CD11b^high^CD45^+^ cells were found in all IRF8KO brains examined (Fig. S2A, S2B). The former subset expressed microglial markers, CX3CR1 and TMEM119, although its levels were lower than WT microglia. P3 cells did not express microglia markers nor border-associated macrophage marker CD206 but expressed Ly6C (Fig. S2C, S2D), indicating the presence of an additional cell population. Detailed characterization of P3 cells will be presented elsewhere. We further assessed the presence of border-associated macrophages (BAM) in our microglia preparations^21, 26^. FACS analysis revealed that cells with CD206 and FOLR2 were less than 1% of the entire brain CD11b^+^CD45^+^ cells prepared from WT and IRF8KO mice (Fig. S2E, S2F).

We performed bulk RNA-seq analyses for adult WT and constitutive IRF8KO microglia and found that the microglia identity genes and CLEAR lysosomal genes were drastically downregulated in IRF8KO cells (Fig.2A, B). The former include *P2ry12, Siglech,* and *Ccr5,* encoding surface proteins and a transcription factor, *Sall1,* while the latter include *Ctsb, Ctsc, Tfeb,* and *Lamp1* involved in waste clearance (Fig. 2B right)^11, 27^. Conversely, interferon-related genes and disease-associated microglia (DAM) like genes, such as *Apoe,* were strongly upregulated in IRF8KO cells^11,28, 29^ (Fig. 2B).

**Figure 2.**
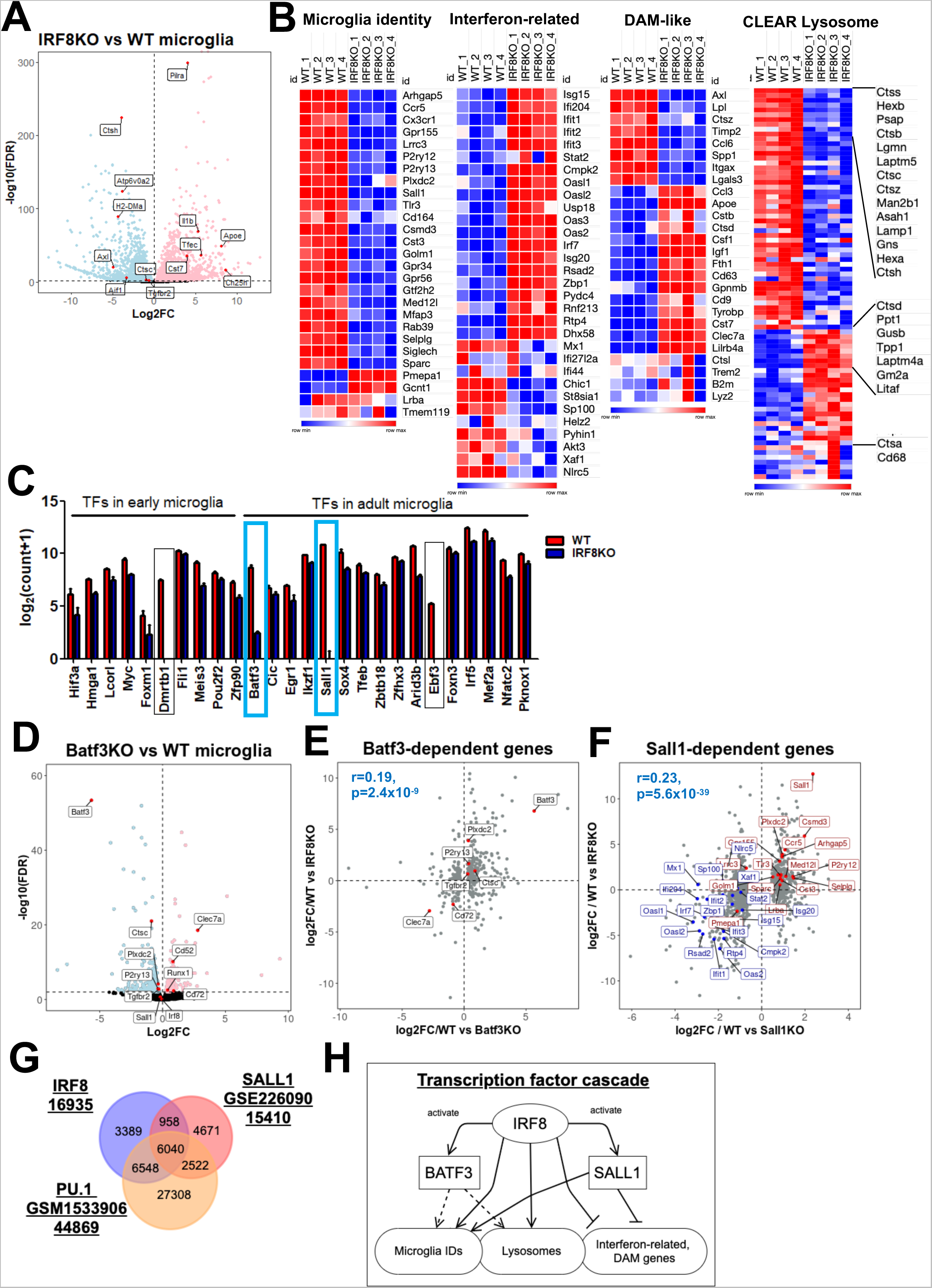
Identification of IRF8-driven transcription factor cascades in shaping microglia-specific transcriptional programs. A. Volcano plot for RNA-seq comparing WT and IRF8KO microglia. Genes upregulated in IRF8KO microglia are indicated in pink (n=2,356), and genes downregulated in IRF8KO cells are in blue (n=2,682, FDR<0.01). Representative genes are shown with a label. B. Heatmaps for indicated gene sets. "Microglia identity" and "Interferon-related" gene sets were referred to as "cluster 6: Microglia" and "Interferon-Related" in the previous publication (ref.11). The DAM-like (ref.29) and CLEAR lysosome gene set were obtained from elsewhere (ref.27). C. Transcription factors downregulated in IRF8KO microglia (FDR<0.01 cut-off) are shown. A list of transcription factor genes was obtained from the previous publication (ref.13). Genes that show more than four log-scale reductions in IRF8KO cells are boxed, and Sall1 and Batf3 in the blue box were further investigated. D. Volcano plot for RNA-Seq data comparing WT and Batf3KO microglia. The genes upregulated in Batf3KO cells are shown in pink (n=172) and downregulated genes in blue (n=274, FDR<0.01). The representative genes are indicated in the plot. E. Correlation biplot showing that Batf3 dependent genes correlate with those dependent on IRF8. The DEGs from Batf3KO microglia RNA-Seq were analyzed with the Kendall rank correlation test (r=0.19, p=2.4×10^-9^). Microglia identity genes shared between Batf3KO DEGs and IRF8KO DEGs are labeled red. F. Correlation biplot showing that Sall1 dependent and IRF8 dependent genes overlap. All up and down DEGs from Sall1KO microglia RNA-Seq obtained from the previous publication (ref.25). were analyzed with the Kendall rank correlation test (r=0.23, p=5.6×10^-39^). The microglia identity genes shared between Sall1KO DEGs and IRF8KO DEGs are labeled red, and those of interferon-related genes are blue. G. IRF8 peaks overlapped with SALL1 and PU.1 binding sites. The SALL1 and PU.1 binding profiles were obtained from GSE226090 (ref.32) and GSM1533906 (ref.33), respectively. H. A model illustrating IRF8 dependent transcription cascade. IRF8 induces expression of downstream TFs, BATF3 and SALL1. These TFs, along with IRF8, then activate indicated target genes. This dual cascade would cover all four gene sets, microglia identity, interferon related, DAM-like, and CLEAR lysosomal genes.

Since IRF8 is known to activate transcription factors to promote myeloid cell development, we examined levels of transcription factors expressed in postnatal microglia^13, 18, 30^. Among them, *Batf3* and *Sall1* were markedly downregulated in IRF8KO cells (Fig. 2C, blue square). BATF3, a basic leucine zipper motif (bZIP) family of a transcription factor, is known to be critical for generating a dendritic cell subset^31^. Because the role of BATF3 in microglial transcriptome remained unelucidated, we next analyzed RNA-Seq in Batf3KO and WT microglia (Fig. 2D). As anticipated, genes downregulated in Batf3KO microglia, such as *P2ry13* and *Ctsc* were also downregulated in IRF8KO cells, indicating that IRF8 and BATF3 control an overlapping set of target genes (Fig. 2E, Table S1 for all overlapped genes). SALL1 is a transcription factor of the Spalt family, expressed selectively in microglia. Buttgereit et al. reported that SALL1 is required for the acquisition of microglia signature^25^. The correlation biplot also showed a significant overlap between SALL1 dependent genes and IRF8 dependent genes, including microglia identity genes (Fig. 2F, Table S1 for all overlapped genes). Fixsen et al. recently identified the genome-wide Sall1 binding profile in microglia^32^. Peak overlap analysis for the IRF8 binding profile revealed that 6,998 (42%) of IRF8 co-bind to SALL1, most of which also bind to PU.1^32, 33^. Results suggest that IRF8, SALL1, and PU.1 cooperatively regulate the expression of those genes (Fig. 2G, Table S2 for all overlapped peaks and their nearest genes)

While IRF8 enhances the expression of *Batf3* and *Sall1*, they do not control IRF8 expression (Fig. 2D). These data indicate that IRF8 acts as an upstream factor in the transcription cascades and activates *Sall1* and *Batf3* expression, which in turn activates downstream target genes (Fig. 2H). We found that a complete microglia transcriptional program is realized after P14, suggesting that epigenome establishment precedes gene expression (Fig. S2G).

### IRF8 configures microglia specific enhancers

To further characterize the chromatin environment in which IRF8 binds, we examined histone marks linked to active transcription, particularly those denoted for enhancers, H3K4me1 and H3K27ac^34, 35^. Of 16,935 IRF8 binding sites, about 73% and 66% of IRF8 colocalized with H3K4me1 and H3K27ac, respectively, and 64% of IRF8 binding sites carried both histone marks (Fig. 3A). However, the histone marks colocalizing with IRF8 peaks were much lower in IRF8KO cells than in WT cells, indicating that IRF8 helps set enhancer histone marks (Fig. 3B, S3A). Since the H3K27ac is a surrogate marker for super-enhancers and super-enhancers neighbor many genes critical for defining cell type specificity and function^35, 36^, we examined super-enhancers using the ROSE program^35^. As expected, many IRF8-dependent microglial identity genes were neighbored by super-enhancers, which totaled 837. Genes neighboring super-enhancers included *Sall1, Siglech,* and *Cst3* (Fig. 3C). Indeed, 735 of 837 (87.8%) super-enhancers were bound by IRF8 (Fig. 3D). While most H3K27ac^high^ regions were populated by IRF8 binding in adult microglia, this was not the case in early postnatal P9 microglia and peritoneal macrophages (Fig. 3E). In addition to super-enhancers, IRF8 was abundantly enriched in smaller enhancers found in specific intergenic and intronic regions overlapping with H3K4me1 (Fig. S3C-G). IGV tracks for the *Sall1* and *Batf3* genes showed that clusters of IRF8 binding coincide with those of H3K27ac and H3K4me1 (Fig. 3F). These results indicate that IRF8 binding guides the formation of enhancers unique to microglia during postnatal development.

**Figure 3.**
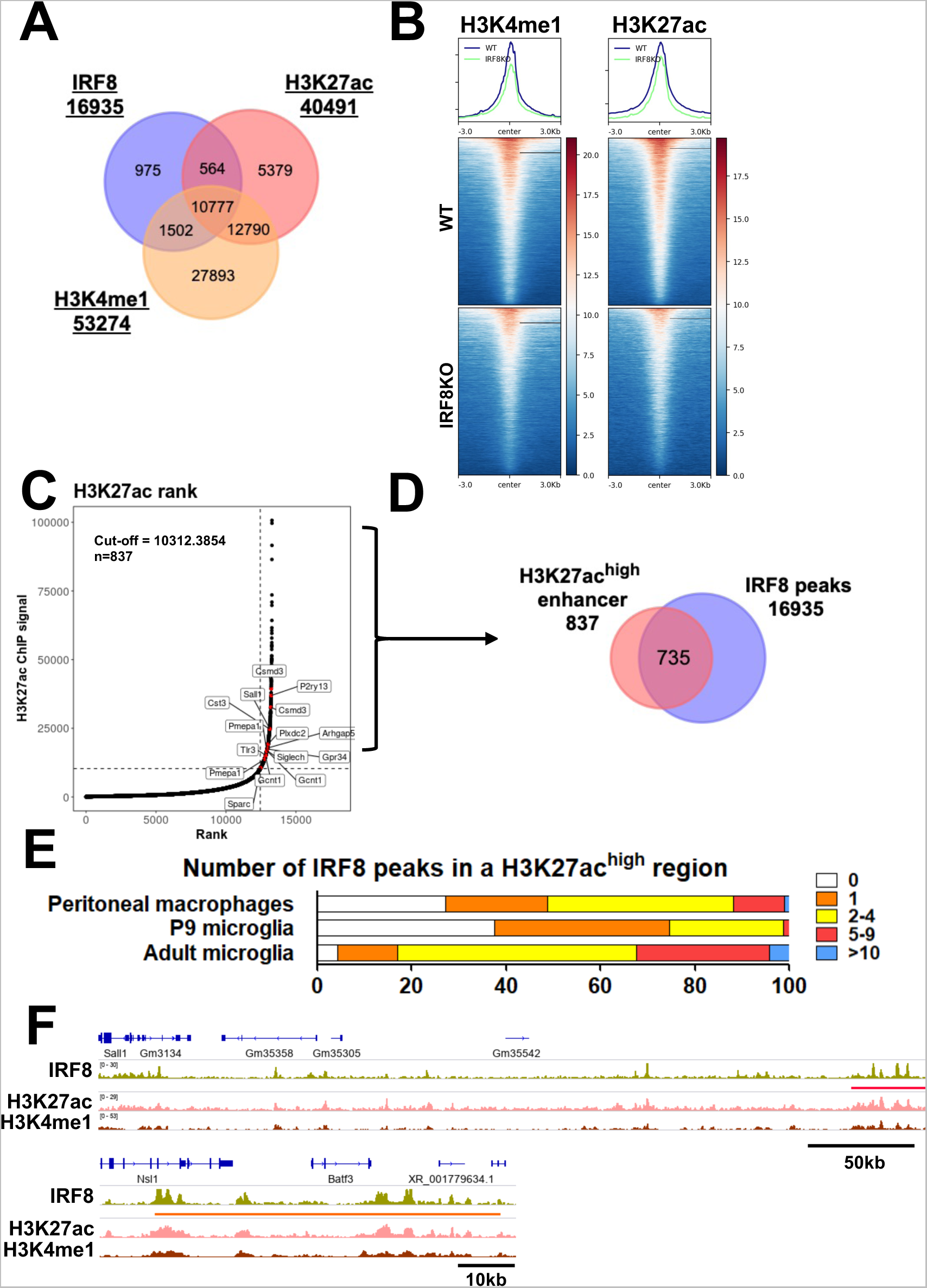
Adult IRF8 is an active enhancer that establishes an enhancer landscape in adult microglia. A. A Venn diagram showing the overlap among IRF8 peaks, H3K4me1, and H3K27ac marks. B. Heatmaps showing H3K27ac (left) and H3K4me1 (right) signal intensity centered on IRF8 peaks ± 3.0kb in adult WT and IRF8KO microglia. Shown above heatmaps are average peak intensities. C. H3K27ac ranked super enhancers in adult WT microglia. IRF8 dependent genes neighboring super enhancers are indicated in red. D. Venn Diagram depicting the overlap of H3K27ac super enhancers and IRF8 peaks in adult WT microglia. E. IRF8 peak counts of adult microglia, P9 microglia, and peritoneal macrophages in the H3K27achigh regions. F. Representative IGVs visualizing the distribution of IRF8 peaks, H3K4me1, and H3K27ac marks near/at Sall1 (top) and Batf3 genes (bottom). An H3K27ac^high^ region is marked by a red bar. IRF8 is bound densely to the Batf3 regulatory region (orange bar), distinct from super enhancers. See also Fig.S3C-G for further description.

### IRF8 confers chromatin accessibility on postnatal microglia and sets DNA methylation patterns

To investigate genomic sites accessible for transcription and whether IRF8 influences the process, we performed an assay for transposase-accessible chromatin (ATAC) sequencing^37^. Differential accessibility analysis revealed that 10,514 peaks were lost in IRF8KO microglia while gaining 5,926 new peaks (Fig. 4A). As anticipated, gained and lost peaks were mostly localized to intergenic regions and introns (Fig. 4B). *De novo* motif analysis showed that lost peaks were enriched with the ETS/ISRE composite motifs (Fig. 4C, left). Gained peaks were enriched with motifs for PU.1, C/EBP, and Smad2/3 binding sites (Fig. 4C, right). Further, ATAC peaks lost in IRF8KO cells were enriched with WT-specific H3K4me1 and H3K27ac marks. On the other hand, IRF8KO-specific H3K4me1 marks were found in gained ATAC peaks (Fig. S3A, 4D). Besides, IRF8KO cells had extra PU.1 binding sites absent in WT microglia (Fig. S3B). Comparison of ATAC-seq peaks and RNA-seq profiles found at lost ATAC-seq peaks in IRF8KO cells were enriched in genes downregulated in IRF8KO cells, such as microglia identity genes (Fig. 4E, in red). Whereas gained peaks were enriched in genes upregulated in IRF8KO cells (Fig. 4E, in purple). IGV tracks showed multiple ATAC peaks on and near *Sall1* and *Batf3* genes in WT cells but few in IRF8KO cells (Fig. 4I). Further, in WT cells, H3K27ac and H3K4me1 marks were abundantly present at these ATAC-seq sites but reduced in IRF8KO cells (Fig. 4I). PU.1 binding sites also revealed a correlation with ATAC-seq peaks (Fig. 4I). However, *Apoe*, expressed in IRF8KO cells but not WT cells showed multiple ATAC-seq peaks in IRF8KO cells, not evident in WT cells. Together, these data indicate that IRF8 sets chromatin accessible sites in microglia, thereby establishing a microglia specific transcriptional program.

**Figure 4.**
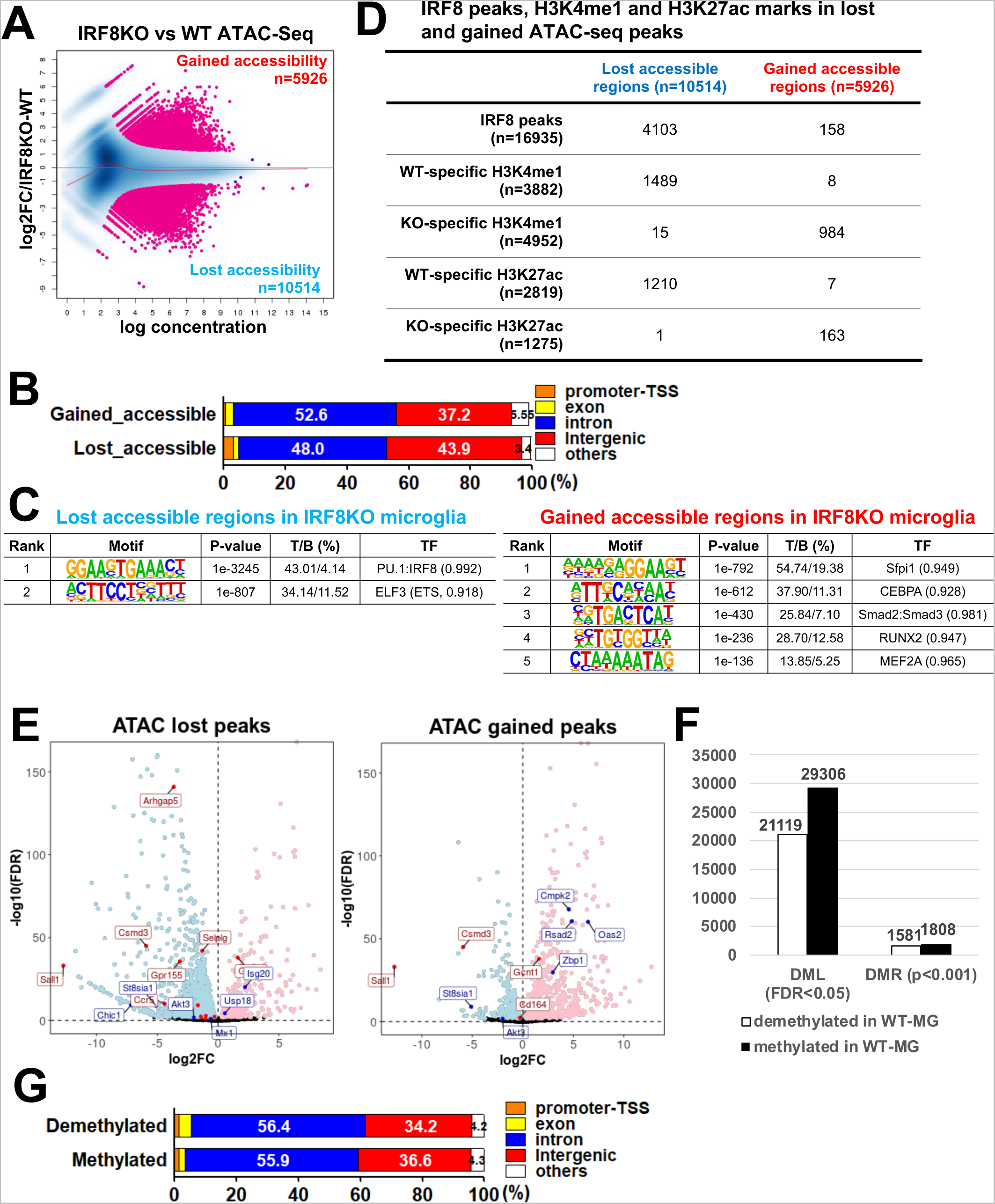
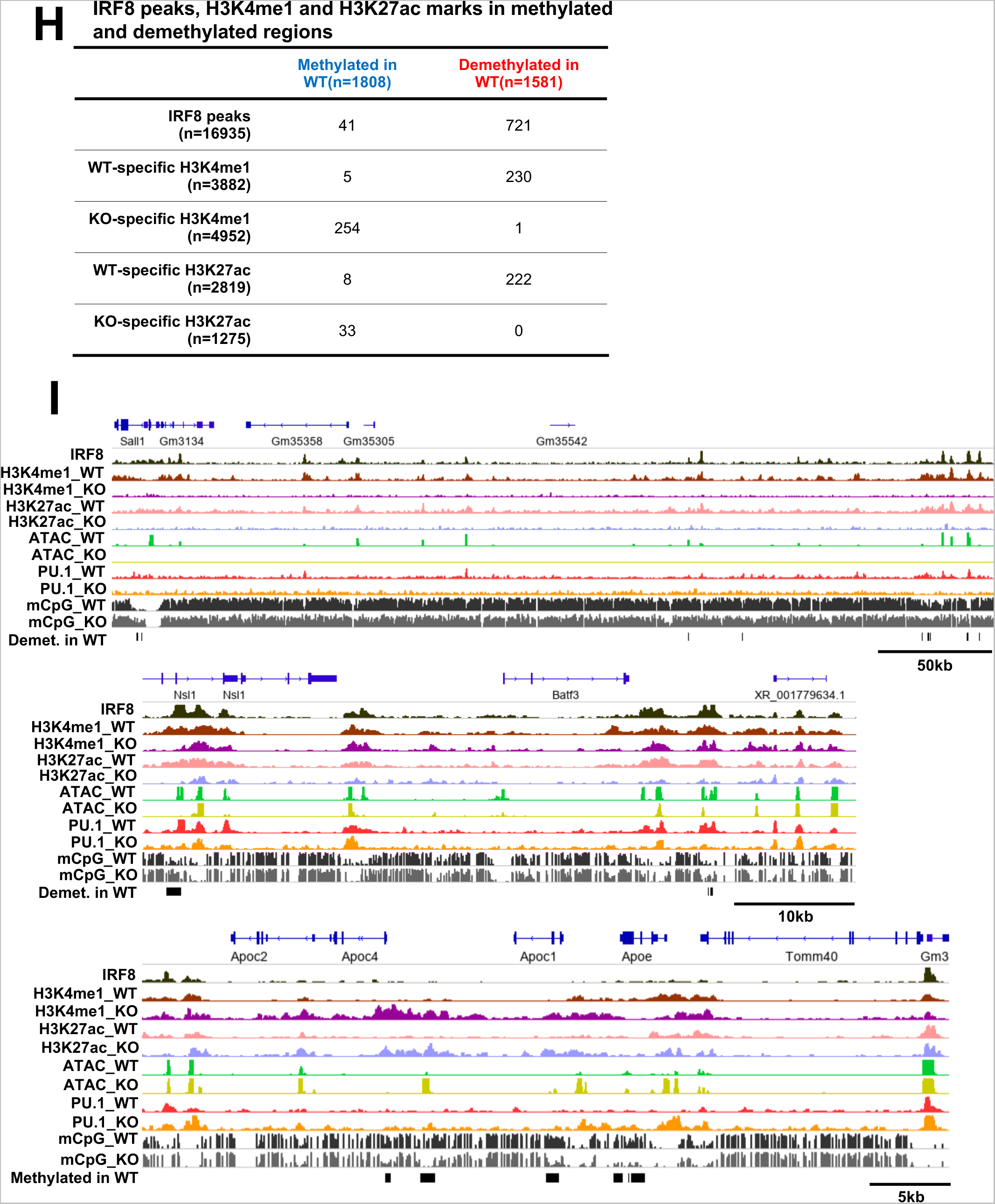
IRF8 confers chromatin accessibility on postnatal microglia and sets DNA methylation patterns. A. MA plot showing ATAC-seq signals comparing WT and IRF8KO microglia (n=2). The differentially accessible regions (FDR<0.05) are colored in pink. The number of gained or lost accessible regions in IRF8KO microglia was shown at the corresponding corner of the area. B. The genome-wide distribution of gained and lost ATAC-seq peaks as determined by HOMER. C. HOMER *de novo* motif analysis for gained and lost ATAC-seq peaks. The top 4 motifs are shown. D. The number of IRF8, H3K27ac, and H3K4me1 peaks (WT or IRF8KO specific) overlapped with lost or gained ATAC-seq peaks. See Fig.S3A for the WT or IRF8KO specific peak identification. E. Volcano plot showing a correlation between ATAC-seq peaks and RNA-seq data. Microglia identity genes (representative "down" genes) and interferon-related genes (representative "up" genes) are labeled red and blue, respectively. The nearest genes for gained or lost ATAC-seq peaks correlate with genes up or down-regulated in IRF8KO microglia. F. The number of differentially methylated single nucleotide loci (DML; FDR<0.05) and the differential regions (DMR; p<0.001). Numbers in the graph are differentially methylated and demethylated loci/regions in WT cells. G. Genome-wide distribution of the differentially demethylated (top) and methylated (bottom) regions identified with HOMER. H. Numbers of IRF8, H3K27ac, and H3K4me1 (WT or IRF8KO specific) peaks in differentially methylated or demethylated regions. I. Representative IGVs visualizing IRF8. H3K4me1, H3K27ac, ATAC-seq, PU.1 peaks and methylated DNA (CpG) status on Sall1 (top), Batf3 (middle), and Apoe (bottom) in WT and IRF8KO cells. Methylated and demethylated regions are indicated by black bars.

To determine whether chromatin accessibility changes during the postnatal stage, we further analyzed ATAC-seq from P9, P14, and adult microglia. ATAC-seq peak numbers increased from P9 to P14, and P14 to adult (Fig. S4A left). Many of these peaks overlapped with IRF8 peaks (Fig S4A right). Consistent with this, *de novo* motif analysis revealed that ETS/ISRE composite elements were enriched within these peaks (Fig. S4A right). In contrast, the Batf3 motif was found during the transition from P14 to adult (Fig. S4B). These data indicate that chromatin accessibility in microglia increases during the postnatal stage in an IRF8-dependent manner. We presented a list of the other transcription factors that may co-bind to IRF8 during the postnatal stages in Table S3. That microglia gene expression increases substantively only after adulthood (Fig S2G), while ATAC-seq peaks increase earlier, supports the view that the formation of epigenome structure precedes that of transcriptional programs^38^.

DNA methylation influences the epigenetic landscape in various somatic cells^39^. In some cells, DNA methylation is linked to gene silencing^39^. However, DNA methylation status has not been deciphered fully for microglia. To determine global DNA methylation sites in microglia and the role of IRF8 in affecting DNA methylation, we performed whole genome bisulfite sequencing. We identified over 20,000 methylated loci (DML) that were differentially regulated in WT and IRF8KO microglia (Fig. 4F)^40, 41^. In addition, 1,808 DNA methylated regions (DMR) and 1,581 demethylated regions were differentially regulated in WT and IRF8KO cells, indicating that IRF8 takes part in DNA methylation in microglia (Fig. 4F). Both methylated and demethylated regions were in intergenic regions and introns, away from the proximal promoter and TSS (Fig. 4G). We found that IRF8 binding sites were located mostly in demethylated regions in WT cells (Fig. 4H, top). Further, methylated regions were enriched with the H3K4me1 and H3K27ac marks present in IRF8KO cells, while demethylated regions were enriched with those marks in WT cells (Fig. 4H). Correlation biplot analysis found that demethylated regions in WT cells correlated with ATAC-seq peaks present in WT cells, while methylated regions correlated with ATAC-seq peaks found in IRF8KO cells (Fig. S4C). These data indicate that demethylated regions were accessible to transcription factors, while methylated sites were not. Consistent with this idea, GO analysis revealed that demethylated regions were enriched with terms such as regulation of immune responses and leukocyte activity (Fig. S4D). In IGV tracks, individual methylated and demethylated sites are shown for *Sall1*, *Batf3*, and *Apoe* (Fig. 4I, bottom two lanes). Sall1 and Batf3 were enriched with demethylated sites in WT cells, but not IRF8KO cells. Conversely, Apoe exhibited many methylated sites in WT cells but demethylated sites in IRF8KO cells. Together, our results indicate that the regulation of DNA methylation is an integral mechanism for setting epigenomic landscape in microglia, which IRF8 plays a critical role.

### Single-cell RNA-seq analysis reveals different clusters for WT and IRF9KO microglia

Next, we performed single-cell RNA-Seq (scRNA-Seq) analysis to assess transcriptome diversity. After filtration, 3,504 WT and 5,303 IRF8KO cells expressing a total of 15,376 genes were selected. Leiden clustering denoted three WT clusters, seven IRF8KO clusters, and a "Common" cluster consisting of WT and IRF8KO cells (Fig. 5A and B). The SigEMD pipeline identified 6,149 DEGs between WT and IRF8KO cells, which covered 62% (3016/4848) of DEGs found in bulk RNA-seq analysis, lending credence to our data (Fig. S5A). Microglia identity gene expression was higher in all WT clusters than in all IRF8KO clusters (Fig. 5C, left). Population heterogeneity was evident within IRF8KO cells, in that ISGs were exclusively found in Cluster 4 (Fig. 5C, middle). In line with this, GO analysis of this cluster ranked interferon-related terms as the top category (Fig. S5B). DAM-like genes were enriched in IRF8KO Cluster 1, although not exclusive (Fig 5C, middle). However, CLEAR lysosomal gene expression was uniformly decreased in all IRF8KO cells (Fig. 5C, right). Interestingly, IRF8KO Cluster 6 did not express microglia identity genes, nor other genes expressed in IRF8KO cells. Instead it expressed *Ly6c2*, encoding Ly6C (Fig. 5C, S5C). Thus, this cluster likely represents the P3 population in IRF8KO brain (Fig. S2C). The Common cluster exhibited unusual features, in that most microglial genes as well as *Ly6c2* were absent in this population (Fig. 5D). This raised the possibility that Common cluster represents a population prior to microglia maturation. We thus applied the PAGA program, which provided a trajectory network (Fig. 5E)^42^. This network illustrated a tight connection with the Common cluster to all WT and IRF8KO clusters except IRF8KO Cluster 6. Another unusual feature of Common cluster was the expression of cell cycle genes such as *Cenpa*, *Ccnb2*, and *Mki67*, a marker for cell proliferation (Fig. 5F). Lastly, Common cluster did not express genes downstream of IRF8, such as Sall1 and Batf3 (Fig.5G). Together, these results may support the idea that Common cluster represents a predecessor of microglia, possessing the capability to develop into functioning microglia and replenish microglia continually for life.

**Figure 5.**
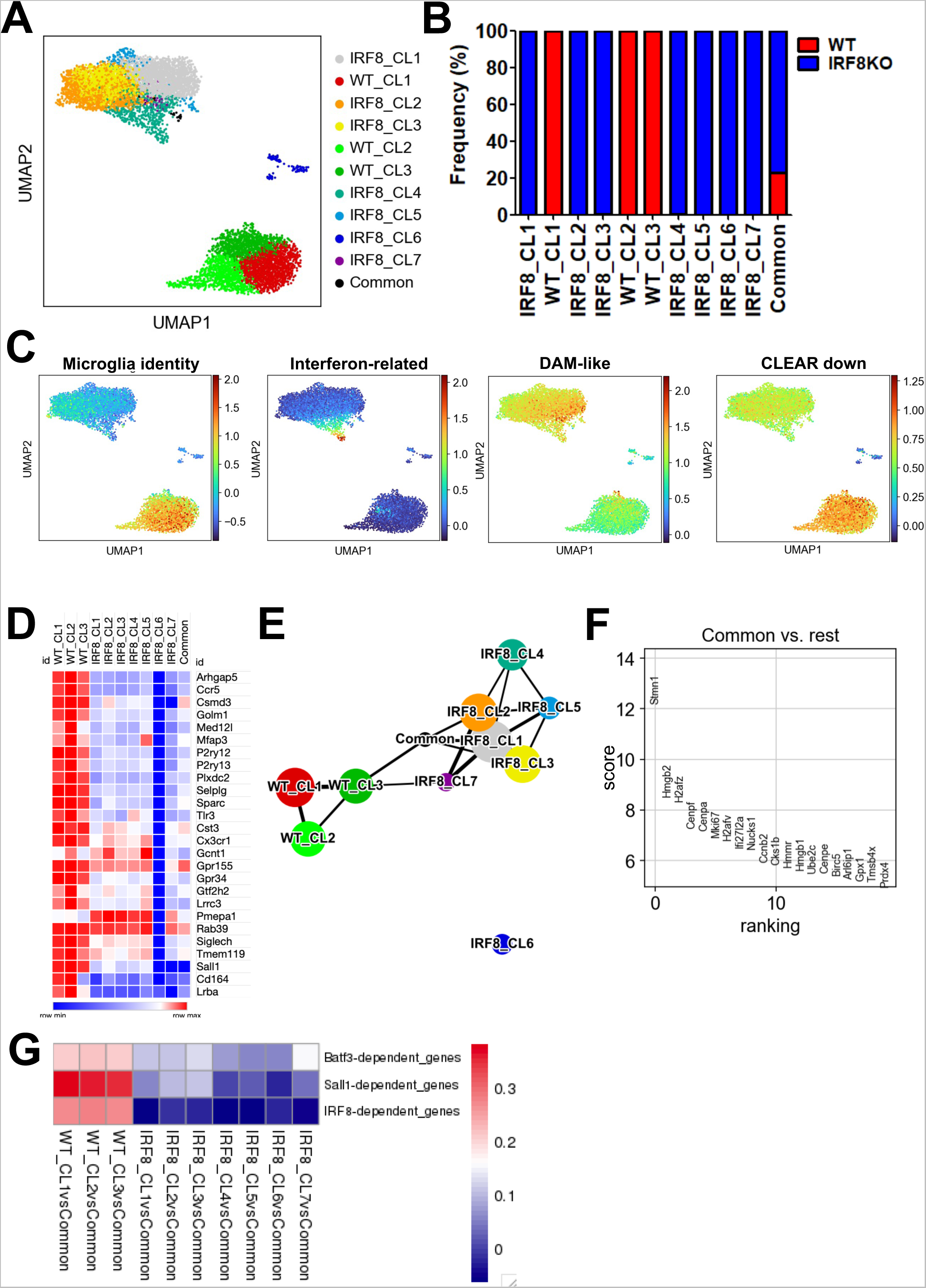
scRNA-seq analysis reveals transcriptome heterogeneity and the presence of a progenitor like population. A. UMAP derived from WT and IRF8KO microglia aggregates. Leiden clusters were colored and labeled as indicated. The data were obtained from the pool of three individual WT and IRF8KO microglia preparations. Percentages of WT and IRF8KO genotypes in indicated clusters. B. UMAP presentation of indicated gene sets (compare with heatmaps in Fig. 2B). C. Gene expression heatmap showing the expression of microglia identity gene set in each WT and IRF8KO cluster. Note that expression of all identity genes is absent in IRF8KO Cluster 6. D. PAGA network depicted with the default parameters. Each node indicates the corresponding Leiden cluster. E. Rank plot aligning the top 20 genes by scoring expression level. The genes in the Common cluster were compared to those of all the other clusters. F. Heatmap displaying the correlation coefficients between Common cluster and other clusters for IRF8, BATF3, or SALL1 dependent gene sets.

### Postnatal IRF8 expression is required for transcription program in adult microglia

To assess whether IRF8 is required for postnatal microglia, we constructed Irf8^flox/flox^Cx3cr1cre^ERT2^ mice to allow for conditional *Irf8* deletion. The mice also carried a Rosa26-loxP-stop-loxP-EYFP reporter, with which we could identify and isolate Cre-expressing microglia^43^ (Fig. 6A). To our dismay, the efficiency of *Irf8* deletion was very poor when Tamoxifen was administered on 4-week-old mice five consecutive times (Fig. S6A). To improve deletion efficiency, we decided to delete Irf8 in younger mice between P12 and P16 and found a considerable improvement in *Irf8* deletion efficiency (Fig. S6A). The basis of poor deletion efficiency is currently unclear, but it may be due to a shift in Cre accessibility. We first performed bulk RNA-seq for IRF8cKO and WT microglia. There were 334 genes downregulated in IRF8cKO cells, while 546 genes were upregulated in IRF8cKO cells (Fig. S6B). Many up and down-regulated genes were included in those found in constitutive IRF8KO cells (Fig S6C). Namely, many microglia identity genes and CLEAR genes were down-regulated, while DAM-like genes and Interferon-related genes were upregulated in IRF8cKO cells, indicating that the same gene sets were affected by constitutive and conditional IRF8 deletion (Fig S6D). The results suggested that IRF8 needs to be expressed in postnatal microglia to maintain the distinctive transcriptome program. Considering that *Irf8* deletion was partial, leaving uncertainty of YFP-based selection, we then performed single-cell multiome in the 10x Genomics platform, which provides scRNA-seq and scATAC-seq data for the same set of cells, enabling us to assess the relation between transcriptome profile and epigenetic landscape. We analyzed 14,319 nuclei after filtering low-quality data. For scRNA-seq data, RNA-guided clustering showed 3 WT and 3 IRF8cKO clusters (Fig. 6B, 6C). There were four additional small clusters of WT and IRF8cKO cells, termed Mixed clusters (Fig. 6B, 6C). As anticipated of incomplete Irf8 gene deletion, some cells from IRF8cKO mice were found within WT clusters, estimating that the deletion efficiency was 85% (Fig.6B, right). Microglia identity and CLEAR genes were downregulated in IRF8cKO clusters, revealing slight heterogeneity (Fig. 6D). DAM-like and Interferon-related genes upregulated in IRF8cKO cells also displayed cluster heterogeneity (Fig 6D). Notably, an equivalent of Common cluster observed in IRF8KO microglia was not evident in the IRF8cKO model. Although the basis of the difference is uncertain, it may be due to the difference in the sample preparation steps, such as permeabilization. Incidentally, the identity of minor Mixed clusters in this analysis is currently unclear.

**Figure 6.**
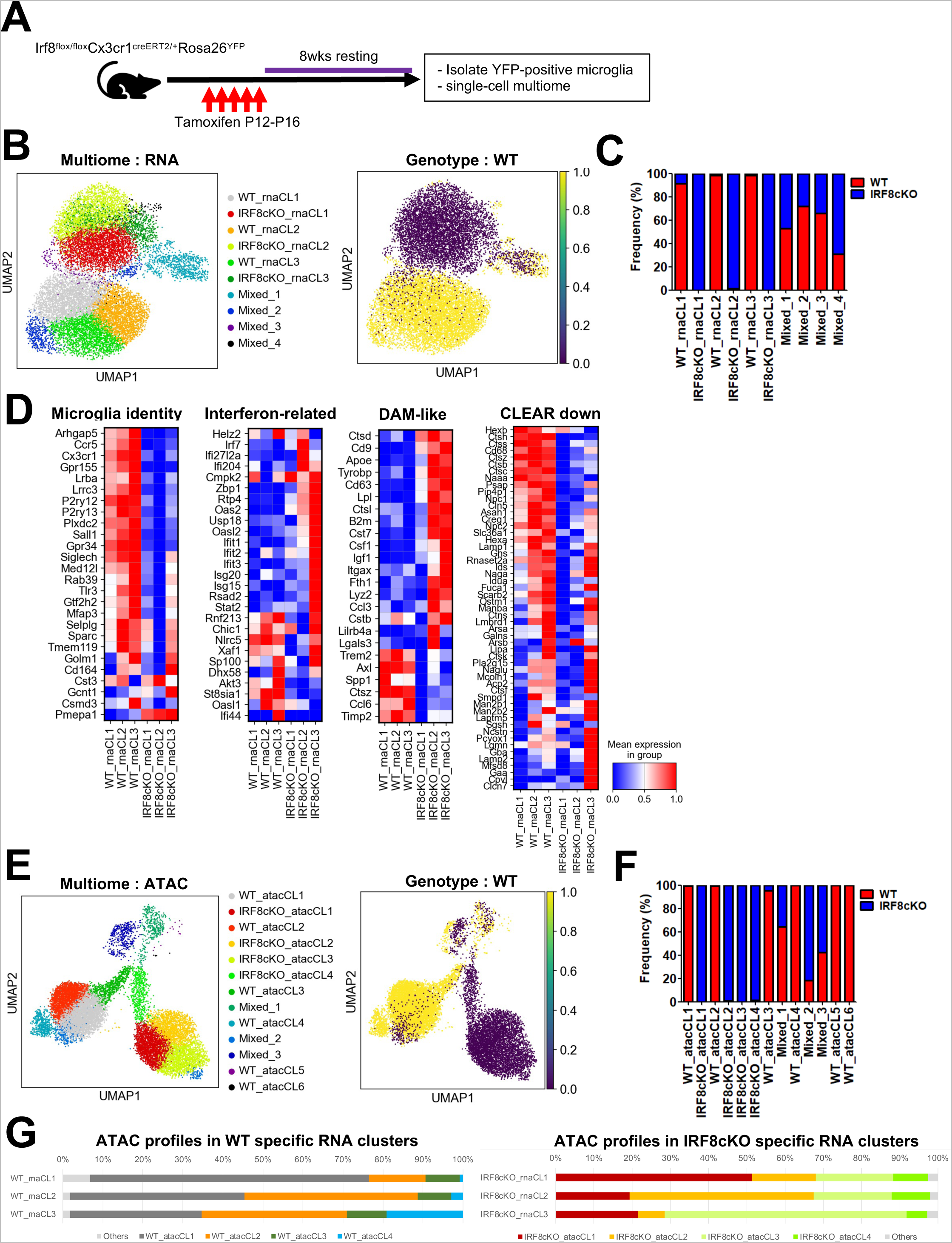
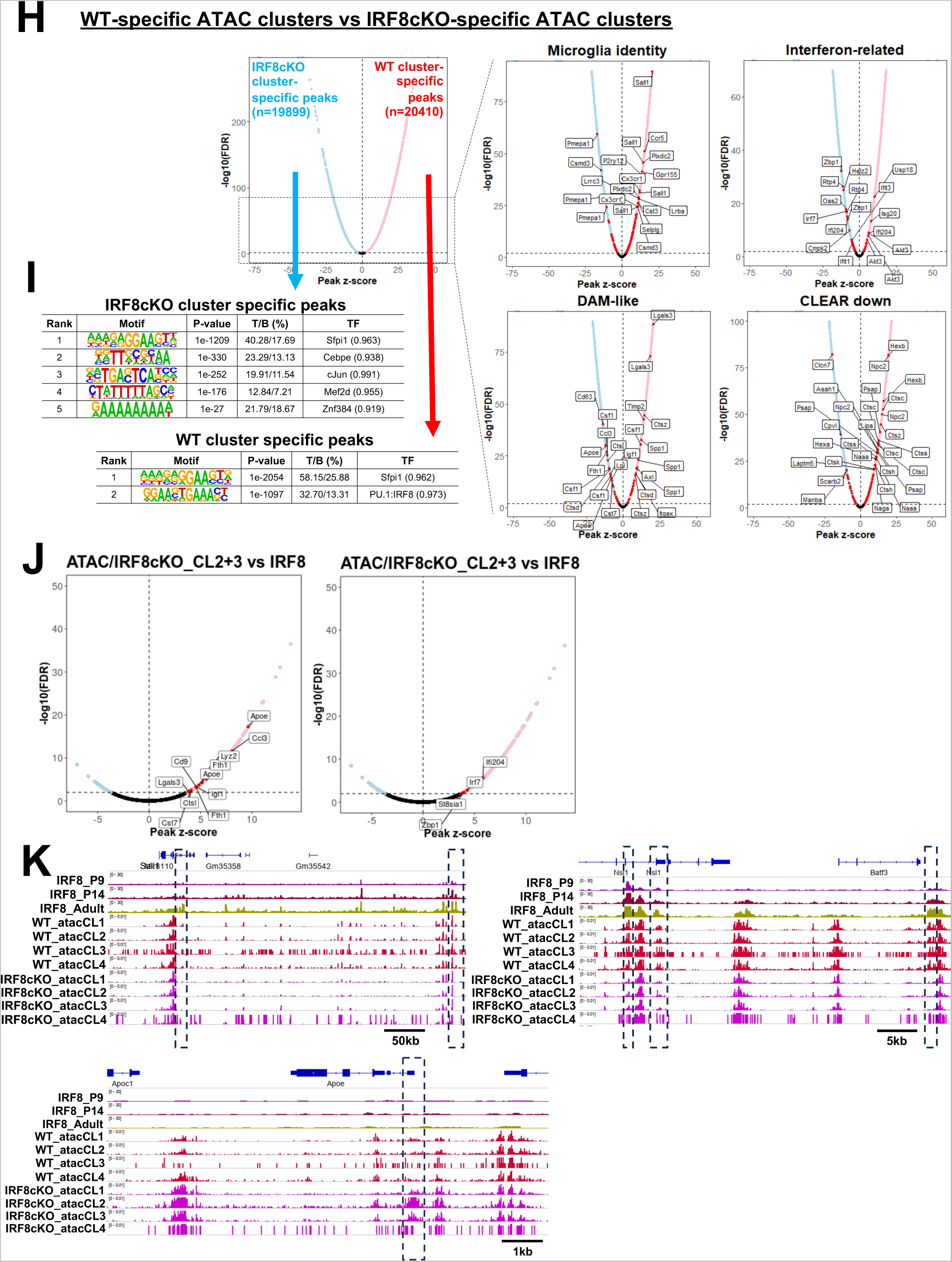
Postnatal deletion of *Irf8* causes gene expression and chromatin accessibility changes analogous to constitutive IRF8KO microglia. A. The strategy for conditional Irf8 deletion in postnatal microglia. P12 Irf8^flox/flox^Cx3cr1Cre^ERT2/+^Rosa26^YFP^ mice were injected with Tamoxifen 5 times, followed by eight weeks of rest. Microglia in adults were isolated and used for bulk RNA-seq and single-cell multiome analysis. B. RNA-guided Leiden clusters on UMAP displaying single-cell multiome sequencing data obtained from WT and IRF8cKO microglia (left). The cells from WT mice were colored yellow in the right panel. C. Frequency of WT and IRF8cKO cells in indicated RNA-guided clusters. D. Heatmap presentation of indicated gene sets (compare with heatmaps in Fig. 2B). E. ATAC-guided Leiden clusters of WT and IRF8cKO microglia, unsupervised with scRNA-seq data (left). The cells from WT mice were colored yellow in the right panel. F. Frequency of WT and IRF8cKO cells in indicated ATAC-guided clusters. G. Percentages occupied by the indicated ATAC-guided cluster cells in WT and IRF8cKO-specific RNA-guided clusters. H. Volcano plot displaying differentially accessible regions. The more accessible regions in WT-specific clusters (WT_atacCL1-4) were colored in pink, whereas those in IRF8cKO-specific clusters (IRF8cKO_atacCL1-4) were in blue. The set of genes presented in Fig.2B was labeled in the four right panels as the nearest genes from peaks, respectively. I. *De novo* motif analysis for differentially accessible regions presented in Fig. 6H J. Volcano plot displaying differentially accessible regions comparing IRF8cKO ATAC cluster 1 and IRF8 ATAC cluster 2+3. DAM-like (left) and Interferon-related genes (right) presented in Fig. 2B were labeled, respectively. K. Representative IGVs of *Sall1*, *Batf3,* and *Apoe* regions. The differential accessible regions were bracketed. IRF8cKO microglia clusters exhibit heterogeneity in the expression of DAM-like and interferon-related genes, which is evident in the *Apoe* bracketed region.

Next, we analyzed the ATAC-seq data to assess chromatin accessibility for the same set of cells. The clustering without reference to transcriptome exhibited six WT clusters, four IRF8cKO clusters, and three mixed clusters containing WT and IRF8cKO cells (Fig. 6E, 6F). Like scRNA-seq analysis, IRF8cKO cells contained WT ATAC clusters (Fig. 6E, right), indicating that IRF8cKO cells that did not delete *Irf8* maintained WT chromatin accessibility. Those scATAC-seq clusters showed expected correspondence with those of RNA-seq clusters, e.g., RNA-seq Cluster 1 in WT cells corresponded to WT ATAC-seq Cluster 1 as evidenced by more than 60% match (Fig 6G). Likewise, IRF8cKO RNA-seq Cluster 1 matched IRF8cKO ATAC-seq Cluster 1 by more than 50%. Nevertheless, considerable intermixing was evident, e.g., IRF8cKO RNA-seq Cluster 2 consisted of roughly 45% ATAC-seq Cluster 2, 40% ATAC-seq Cluster 1, and 15% ATAC-seq Cluster 3 cells (Fig 6G). These results suggest that each RNA-seq cluster carries a corresponding epigenome structure, although there is certain interchangeability.

There were a total of 19,899 lost peaks and 20,410 gained peaks in IRF8cKO cells (Fig. 6H). *De novo* motif analysis showed that lost peaks are enriched with ETS/ISRE composite elements, whereas gained peaks were enriched with PU.1 and C/EBP elements, consistent with bulk ATAC-seq data in IRF8KO cells (Fig. 6I). Lost peaks found in all three IRF8cKO clusters included the majority of microglia identity and CLEAR genes, although some heterogeneity was evident (Fig, 6G, right). Conversely, DAM-like and Interferon-related genes were shown to be in the gain of ATAC peaks in IRF8cKO clusters (Fig 6H, right). This correlation supports a direct mechanistic link between chromatin accessibility and transcription operating at a single-cell level. Additionally, scATAC-seq clusters 2 and 3 in IRF8cKO correlated with the upregulation of DAM-like and Interferon-related genes that dictate heterogeneity of IRF8cKO RNA-seq clusters (Fig. 6J).

IGV profiles for scATAC-seq peaks for *Sall1* and *Batf3* regions show prominent peaks in all WT clusters but not in IRF8cKO clusters (Fig 6K, top). On the other hand, *Apoe,* one of the upregulated genes in IRF8cKO clusters 2 and 3, exhibited a gain of ATAC peaks in IRF8cKO clusters 2 and 3 but not WT clusters (Fig 6K, bottom). Together, maintaining transcriptional program and chromatin accessibility in adult microglia requires continuous IRF8 expression.

### Irf8 deletion ameliorates AD pathology in the mouse model

IRF8 has been shown to promote neuroinflammation and is implicated in AD pathogenesis in mice^19, 23,44, 45^. To further address the role of IRF8 in AD, we first compared microglia transcriptome profiles of WT and 5xFAD mice at 9 months of age^46^. We found 699 genes upregulated in 5xFAD microglia over WT cells, which we regarded as the 5xFAD-associated gene set. This gene set contained many DAM-like genes, such as *Apoe*, *Tyrobp*, and *Cst7* (Fig. S7A)^10,29^. Interestingly, many DAM-like genes were upregulated in IRF8KO microglia without the 5xFAD background (Fig. 7A)^28, 45, 47^. Further, microglia from 5xFAD/IRF8KO mice showed essentially the same expression pattern as IRF8KO microglia (Fig. 7A). Grubman et al. reported that 5xFAD microglia *in vivo* have two distinct populations, one with engulfed Aβ, and the other without, which could be distinguished by staining with a brain-permeable Aβ probe, MethoxyX04 (X04)^48^. X04^+^ cells, but not X04^-^ cells, expressed DAM-like genes along with interferon-related genes^48^. However, mechanisms of setting transcriptome programs in X04^+^ and X04^-^ microglia have remained unclear. Thus, we examined the chromatin accessibility of X04^+^ and X04^-^ cells isolated from 9-month-old 5xFAD brains (Fig. S7B). While 1-2% of microglia in 5xFAD/WT brains were X04^+^, X04^+^ cells in 5xFAD/IRF8KO brains were much less (<0.1%, Fig. S7C), too few for ATAC-seq experiments. The differential analysis for X04^+^ microglia revealed that 1,650 gained peaks, while 26,882 lost peaks in X04^+^ cells relative to X04^-^ cells (Fig 7B). Motif analysis indicated that lost peaks were enriched with PU.1, and ETS/ISRE composite elements, whereas gained peaks were elements for Osr1 and Rfx3 (Fig. 7C). Gained peak motifs in X04^+^ cells differed from those in IRF8KO microglia. Nearest gene analysis exhibited no association between a set of DAM and Interferon-related genes and gain of ATAC peaks, in contrast to IRF8KO and IRF8cKO microglia (Fig. 7D). These results suggested that 5xFAD microglia assume chromatin accessibility patterns different from that in IRF8KO and IRF8cKO microglia. This was evident in IGV profiles of *Apoe* and *Sall1* genes where ATAC-seq peaks in IRF8KO and X04^+^ cells differed (Fig. 7E) and the heatmaps showing scATAC-seq and bulk ATAC-seq signals near DAM genes (Fig. S7D). These data indicate that 5xFAD microglia have epigenome structures unrelated to those in IRF8KO and IRF8cKO cells.

**Figure 7.**
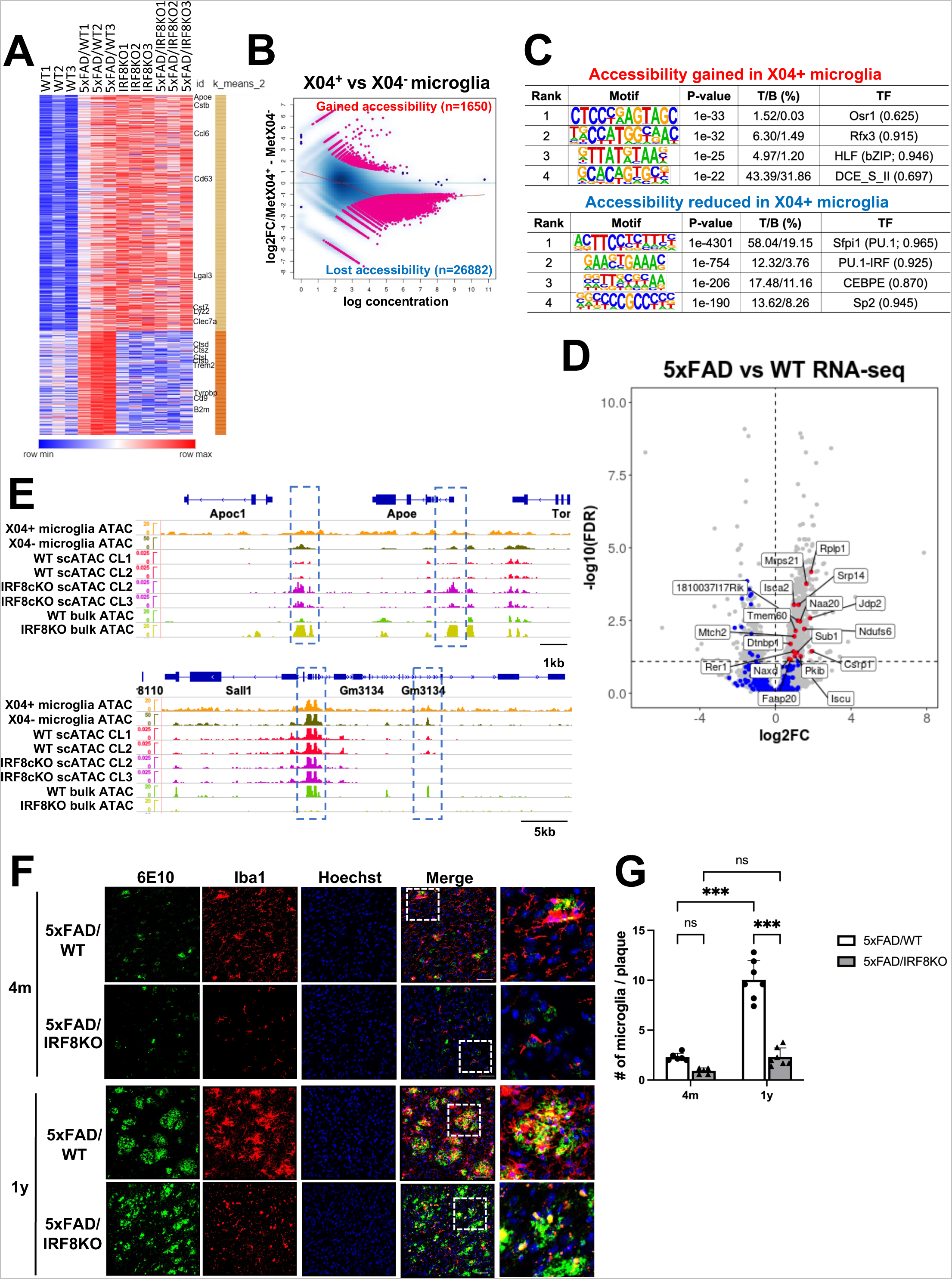
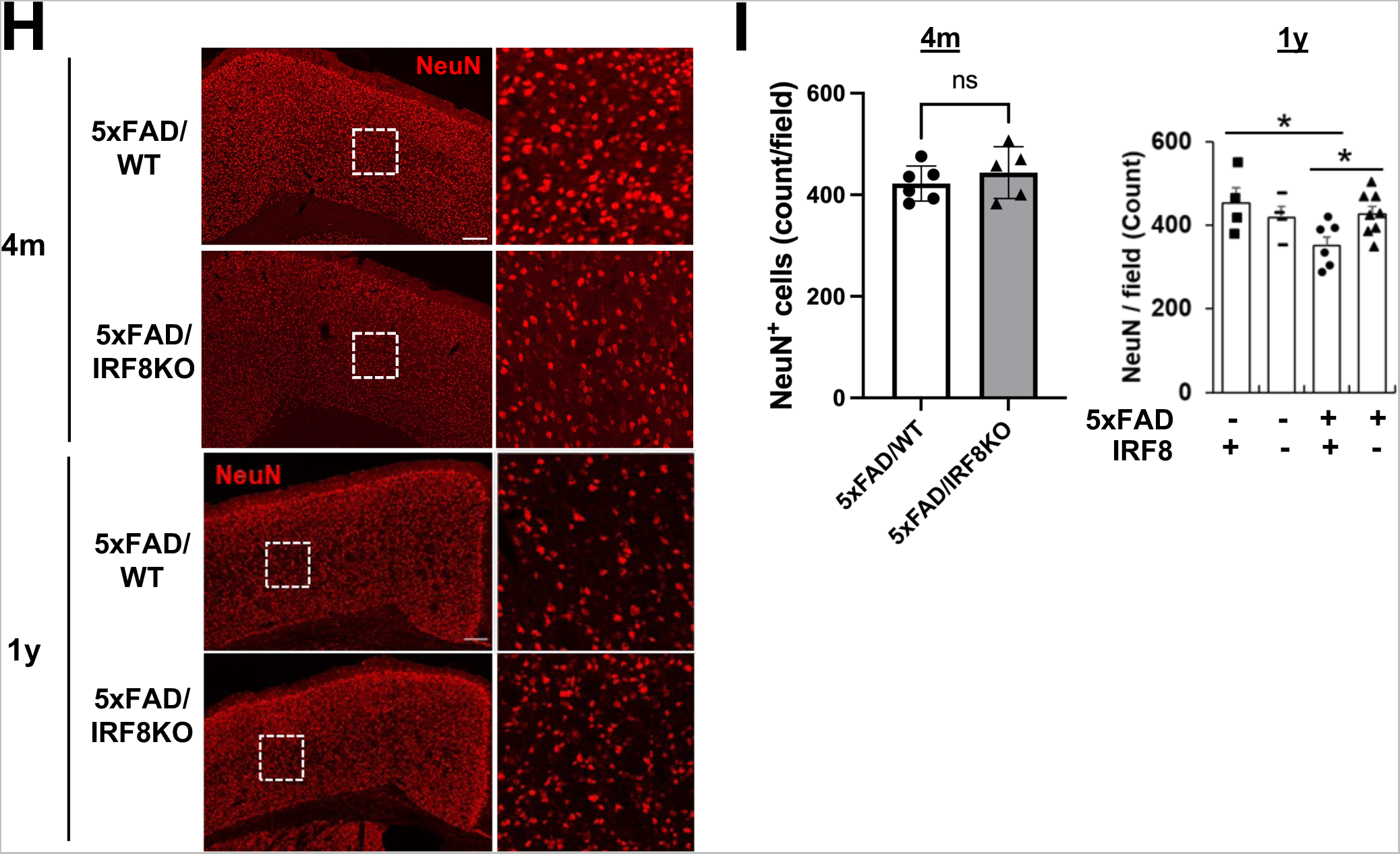
IRF8KO microglia lessens AD pathology in the 5xFAD mouse model, despite spontaneous DAM/NDG gene induction. A. Heatmap showing the 5xFAD associated gene set expression in WT, 5xFAD/WT, IRF8KO, and 5xFAD/IRF8KO microglia. Data are from RNA seq analysis of microglia from nine months old mice. Data were clustered by K-mean methodology (k=2), and representative genes were labeled on the right. B. An M-A plot showing ATAC-seq data comparing MethoxyX04-positive and-negative microglia from 10 months old 5xFAD mouse (n=2). The differentially accessible regions (FDR<0.05) are colored in pink. The number of gained or lost accessible regions in MethoxyX04-positive microglia was shown at the corresponding corner of the area. C. *De novo* motif analysis for differentially accessible regions presented in Fig. 7B D. Volcano plot showing gene expression profiles comparing 5xFAD/WT and non-5xFAD control (same data as Fig.S7A with rescaled). The genes that are nearest to the gained peaks in X04^+^ microglia were plotted in blue. Among them, significantly upregulated genes in 5xFAD microglia (referred to as 5xFAD-associated genes) were highlighted in red and labeled. The gain of ATAC in X04^+^ microglia exhibits some connection with transcriptome in 5xFAD microglia rather than DAM-like genes. E. Representative IGVs of *Apoe* and *Sall1* regions. The X04^+^ microglia do not show an increase of chromatin accessibility at the promoter region of *Apoe* gene (bracketed), whereas IRF8KO or IRF8cKO microglia exhibited a significant increase in that region. For the *Sall1* region, the accessibility was lost entirely in IRF8KO, while some were retained in IRF8cKO and X04^+^ microglia in a cell-type-specific manner (see bracketed regions). F. Representative histology images depicting immunostaining of the cortex regions of 4 months (top) and 1-year-old (bottom), 5xFAD/WT or 5xFAD/IRF8KO brain with 6E10 for Aβ, Iba1 for microglia, and Hoechst33342 for nuclei, respectively, to quantify microglia colocalization with plaques. Merge images on the right illustrate a lack of extensive colocalization in the 5xFAD/IRF8KO brain. On the right are enlarged images from the bracketed regions on the left. Scale bar; 50 µm. G. Quantification of microglia colocalized with plaques in Fig.7F. The data was statistically tested using one-way ANOVA followed by Tukey’s post hoc test (F(3,20)=2.788, p<0.001). Graph bar; mean +/-SEM. H. Representative histology images of NeuN staining in the cortex area of 4 months (top) and a year-old (bottom) 5xFAD/WT or 5xFAD/IRF8KO brain. Scale bar; 200 µm. On the right are enlarged images from the bracketed regions on the left. I. Quantification of NeuN positive cells. Values are the average of five fields per brain. The data was statistically tested using one-way ANOVA followed by Tukey’s post hoc test (F(3,33)=2.570, p<0.001). Graph bar; mean +/-SEM.

**Figure 8.**
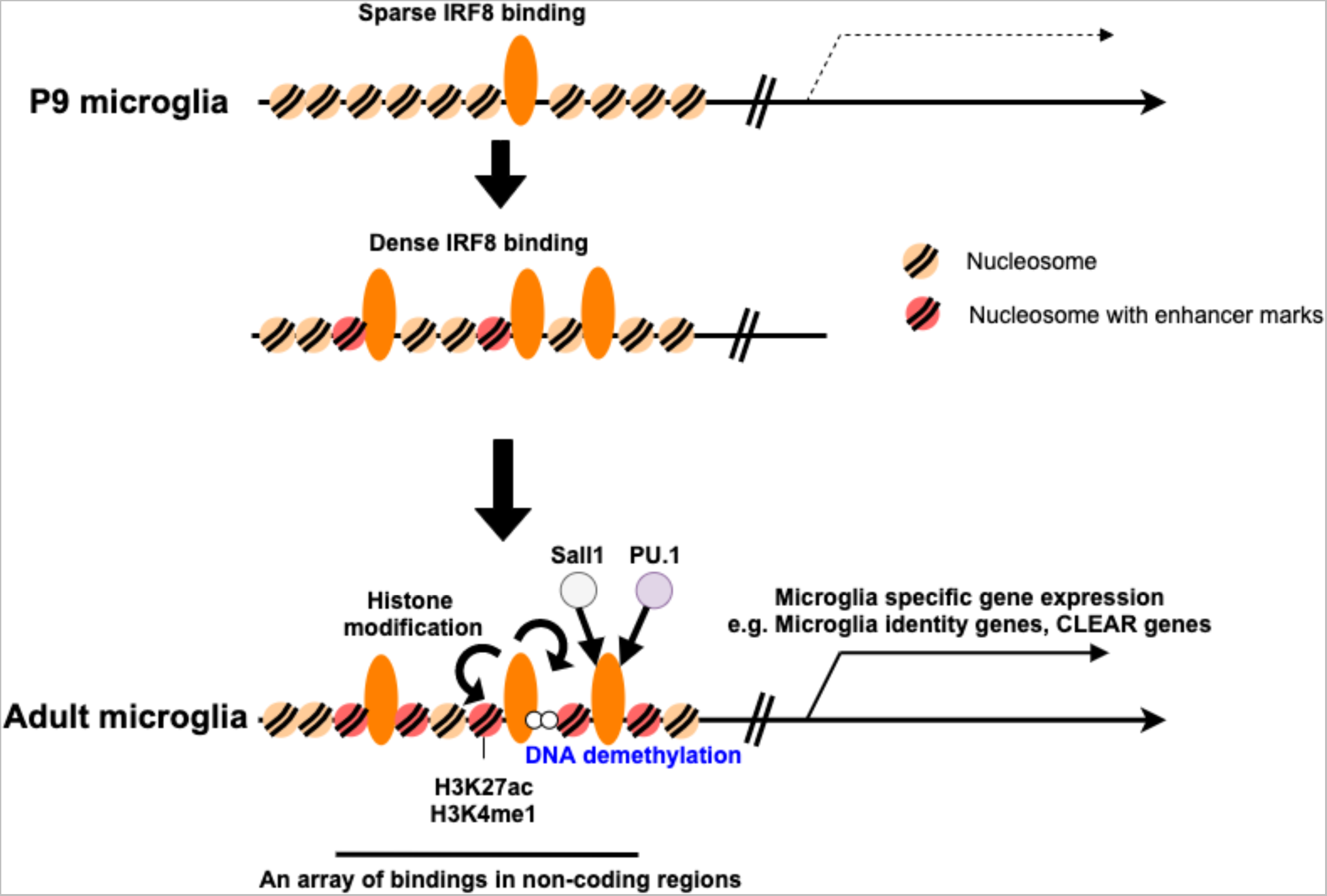
Model for the role of IRF8 in postnatal microglia development At the early postnatal stage (P9 microglia), IRF8 binding to the microglia genome is sparse (Top). After P14 through adulthood, IRF8 binding becomes denser, forming arrays along with histone modifications (Middle). Full IRF8 binding leads to the formation of microglia-specific enhancers along with Sall1 and PU.1, opens relevant regions of chromatin and stably demethylates enhancer regions of DNA (bottom), which, taken together, establishes transcriptional programs in microglia.

We next examined how the loss of IRF8 influences AD pathology. d’Errico et al. recently reported that *Irf8* depletion leads to reduced microglia motility, resulting in reduced spreading of Aβ plaques^44^. However, whether postnatal *Irf8* deletion affects AD pathology and mechanisms by which Aβ plaque spreads within the brain, and the role of IRF8 in the process remained elusive. We performed histological analyses on four-month and one-year-old 5xFAD/WT and 5xFAD/IRF8KO brains for localization of microglia and Aβ in the cortical region (Fig. 7F). The microglia count per plaque was much fewer in 5xFAD/IRF8KO brain compared to 5xFAD/WT brain, both at four months and one year, the extent being greater at one year (Fig. 7F, 7G). These data are in line with the FACS data that 5xFAD/IRF8KO brain has fewer X04^+^ microglia (Fig. S7C). These results indicate that IRF8KO microglia are unable to recognize Aβ plaques^44^. Significantly, IRF8cKO/5xFAD brain microglia were also defective in interacting with Aβ (Fig. S7E). Our data show that postnatal *Irf8* deletion is sufficient to impair microglia’s ability to interact with Aβ.

Additionally, we conducted a study to track the growth of Aβ plaques from 4 months to one year using thioflavin S (ThioS), which detects polymerized Aβ^49^. We found that plaque count and size increased in both 5xFAD/WT and 5xFAD/IRF8KO brains during the period. However, at one year, the size of plagues was much smaller in 5xFAD/IRF8KO brains than in 5xFAD/WT brains. In contrast, the sizes were comparable between 5xFAD/WT and 5xFAD/IRF8KO brains at four months old (Fig. S7F). These data also indicated a defective Aβ plaque growth without IRF8.

Next, we assessed overall neuronal health in 5xFAD/WT and 5xFAD/IRF8KO brains using the NeuN antibody, which detects intact neurons^54^. The NeuN immunoreactivity in 5xFAD/IRF8KO brains was higher than that in the 5xFAD/WT brains at one year, while it was similar at four months (Fig. 7H). These results indicate that the deletion of IRF8 lessens neuronal damage in the 5xFAD brain.

## Discussion

This study aimed to elucidate postnatal microglia development and the role of IRF8 in the process, a question that remained unanswered. We show that IRF8 binds to microglia enhancers in a stepwise fashion during P9-P14 to the adult stage. This coincided with a stepwise increase in chromatin accessibility. However, microglia specific transcriptome expression took place later in adults. Thus, IRF8 binding and setting of open chromatin preceded microglia-specific gene expression, a process consistent with other developmental models^16, 38^.

It was striking that a stepwise increase in IRF8 binding was not due to corresponding increase in IRF8 expression, since IRF8 levels were constant during this period and through adulthood, suggesting an epigenome-based mechanism, yet to be deciphered. Many IRF8 peaks were in microglia specific enhancers neighboring microglia identity and CLEAR lysosomal genes, the former include transcription factors, Sall1 and Batf3, that enhance the expression of these gene sets.

IRF8 deletion led to the loss of these enhancers, resulting in a marked reduction in the expression of these gene sets. IRF8 loss led to the formation of alternative enhancers that were near DAM-like genes (e.g., *Apoe*) and interferon-related genes that were upregulated in IRF8KO and IRF8cKO cells. We further show, by genome-wide bisulfite sequencing, that IRF8 sets DNA methylation patterns specific for microglia, which correlated with ATAC-seq patterns.

scRNA-seq analysis revealed distinct clusters in WT and constitutive IRF8KO microglia, indicating considerable heterogeneity in gene expression patterns, most notable being interferon-related gene expression restricted to one cluster in IRF8KO microglia.

10x Multiome analysis for IRF8cKO microglia was highly informative since we could compare transcriptome profiles and chromatin accessibility patterns for the same set of cells at the single-cell level. Data revealed distinct clusters of IRF8cKO and WT cells. We found marked downregulation of microglia identity gene and CLEAR genes in distinct scRNA-seq clusters, while DAM-like and interferon-related genes were upregulated in the other distinct clusters. scATAC-seq analysis also identified distinct clusters for WT and IRF8cKO cells. A salient finding was that scATAC-seq clusters exhibited good correspondence with scRNA-seq clusters, although some interchangeability was evident. Results suggest that chromatin accessibility defines transcriptome profiles at single-cell level. Certain levels of interchangeability were interesting, as they suggest additional complexity in chromatin accessibility and gene expression. Multiome analysis also enabled us to estimate deletion efficiency in IRF8cKO accurately, which was about 85%. That way, we could remove the contribution of contaminating undeleted cells from our data. We conclude that postnatal IRF8 expression is necessary for microglia to maintain epigenome structure and transcriptional programs for adult microglia.

It has been shown that the microglia from constitutive IRF8KO mice do not spread AD plaques to other regions in the brain, presumably reducing AD pathology^44^. To gain a mechanistic understanding of the role of IRF8 in AD, We examined (a) 5xFAD/IRF8cKO microglia and (b) development of Aβ plaques from early to late stages of the disease, questions that remained unanswered. Remarkably, microglia’s ability to interact with Aβ plaques was lost by constitutive IRF8KO and postnatal *Irf8* deletion, even though deletion was incomplete. Thus, loss of IRF8 impairs microglia’s ability to recognize Aβ without delay. Further, we show that Aβ plaques develop during four months to one year in both number and size and that the absence of IRF8 affects mostly the growth of plaque size. These observations add to our understanding of the role of IRF8 in AD pathology further. Lastly, we addressed the relationship between DAM-like gene expression in IRF8KO and 5xFAD brains. ATAC-seq analysis of X04^+^ 5xFAD and IRF8KO microglia displayed distinct patterns of chromatin accessibility unrelated to each other. Thus, DAM-like gene expression found upon *Irf8* deletion does not seem to have functional relevance to AD pathology in the 5xFAD model.

In summary, this study demonstrates the essential requirement of IRF8 in setting the epigenome structure of postnatal microglia and the role of AD progression.

### Online Methods

#### Mice

B6(Cg)-*Irf8*^tm1.2Hm^/J (IRF8KO), B6.Cg-*Irf8*^tm2.1Hm^/J (Irf8-GFP), B6.Cg-*Irf8*^tm1.1Hm^/J (Irf8flox) and control wild-type C57BL/6J mice were maintained in the NICHD animal facility^23^. B6.129S(C)-*Batf3^tm1Kmm^*/J (Batf3KO), B6.129P2(Cg)-*Cx3cr1*^tm1Litt^/J (Cx3cr1^GFP^), B6.129P2(C)-Cx3cr1^tm2.1^(cre/ERT2)^Jung^/J (Cx3cr1cre^ERT2^), and B6SJL-Tg^(APPSwFlLon, PSEN1*M146L*L286V)6799Vas^/Mmjax (5xFAD) mice were purchased from Jackson Laboratory. Cx3cr1^GFP^ mice were crossed with IRF8KO mice. 5xFAD mice were backcrossed to C57BL/6J strain for at least seven generations and maintained hemizygote. To achieve microglia specific Irf8 depletion (IRF8cKO), we crossed Irf8flox mice with Cx3cr1cre^ERT2^ mice. Mice were given 300µg of Tamoxifen (Sigma) for five consecutive days at P12 or 2mg x 5 days at P28, and the Cre recombinase expression in microglia was confirmed using Rosa26^LSL-EYFP^ reporter mice. For *in vivo* amyloid staining, 2mg/kg Methoxy-X04 (TOCRIS, #4920) in a half DMSO/saline (50%DMSO, 0.9%NaCl, pH12.0) was intraperitoneally injected 3hr before microglia isolation. Female mice were used in this study unless specified, and all animal experiments were performed according to the animal study (ASP#17-044, 20-044, and 23-044) approved by the Animal Care and Use Committees of NICHD, NIH.

### Microglia preparation

Mice were euthanized by asphyxiating with CO_2_ and transcardially perfused with 10 mL of ice-cold PBS. The brain was harvested, chopped with a single-edge razor blade, and ground in ice-cold Hanks-balanced salt solution using a 7ml Dounce homogenizer, then filtered through a 70µm cell strainer. The cells were resuspended in 30% isotonic Percoll and spun down 800xg for 30 min. The floating myelin layer was removed, and the cell pellet was further sorted by FACS AriaIIIµ (BD bioscience) after antibody staining. All reagents and instruments were kept in our 4°C cold room, and all procedures were performed at less than 4°C to avoid extra activation. For FACS cell sorting, the following antibodies were used: CD11b-Brilliant Violet 421 (CD11b-BV421, clone: M1/70, Biolegend), CD45-APC (clone: 30-F11, Invitrogen). The dead cells were gated out with staining of 7-Aminoactinomycin D (7-AAD; BD pharmingen, 1:100). The live cells were counted with a hemocytometer, which was regarded as the 7-AAD negative subset in FACS for estimating the count of each cell population. To efficiently isolate cells, we took a backgating strategy in which all gates for cell sorting were forced to be set where microglia most abundantly existed. The microglial gate was determined with a 10s pilot run before an actual cell sorting. We used CX3CR1-PE (FAB5825P, R&D), Ly6C-APCCy7 (clone: HK1.4, Biolegend), F4/80-PECy7 (clone: BM8, Biolegend), CCR2-BV510 (clone: SA203G11, Biolegend), 1A8-FITC (BD Pharmingen), Tmem119-Alexa Fluor 488 (ab225497, Abcam), CD206-PE/Dazzle594 (clone: C068C2, Biolegend), FRβ-APC (clone:10/FR2, Biolegend), CD3e-PE (clone:145-2C11, Biolegend), NK1.1-PE (clone: PK136, eBioscience), B220-PE (clone: RA3-6B2, Biolegend) antibodies for the expression analysis and reanalyzed the data with FlowJo v.7.6.5 or 10.10.0 software.

### Immunohistochemistry

Mice were anesthetized with 125mg/kg Ketamine and 10mg/kg Xylazine cocktail intraperitoneal injection. The brain was fixed by the transcardial perfusion of 4% PFA/PBS, followed by overnight incubation in an additional 4% PFA/PBS. After removing the fixative, the brain was cryoprotected with a 10-30% sucrose solution. The section was made with 30µm slices. Blocking was achieved with 4% BSA in PBS and 0.3% Triton X-100. Primary antibodies and working dilution are the following: Mouse anti-β-Amyloid antibody (1:500, clone: 6E10, Biolegend), Rabbit anti-NeuN monoclonal antibody (clone: A60, Millipore), and Rabbit anti-Iba1 antibody (Wako). The tissue was then incubated with the following fluorescence-labeled secondary antibodies: Alexa Fluor 488 Goat anti-mouse IgG (H+L) (Invitrogen), Alexa Fluor 568 Goat anti-mouse IgG (H+L) (Invitrogen), Alexa Fluor 633 Goat anti-mouse IgG (H+L) (Invitrogen), Alexa Fluor 633 Goat anti-rabbit IgG (H+L) (Invitrogen). The core-dense amyloid plaques were stained with 0.5% Thioflavin S (Sigma, T1892). The nuclei were stained with Hoechst33342 dye. For each brain section, z-stack images were taken on a Zeiss AiryScan 880 confocal microscope using either a 10x or a 40x air objective, together with the Airyscan system and ZEN lite software (Zeiss). Alexa Fluor 488/Thioflavin S, Alexa Fluor 633 and Alexa Fluor 568, and Hoechst33342 dyes were excited using the 488nm, 633nm, 561nm and 405nm laser lines. The emitted fluorescence was detected using the corresponding channels on the Airyscan detectors. Raw data were further analyzed with ImageJ/Fiji^50^. Maximum intensity projection for the z-stacked images was presented, and the signal from each channel was pseudocolored as specified in the figures. The amyloid burden, plaque size, and count were quantified with five 500 μm² fields randomly chosen from each section. To quantify microglia gathered with a plaque, we randomly picked ten plaques per mouse that were positive for 6E10. We then counted the number of Iba1-positive microglia colocalized with each plaque^44^. The nuclei were stained with Hoechst33342 dye to distinguish Iba1-positive microglia as individual cells. The average microglia count per plaque was then calculated and presented for each mouse^44^. For NeuN-positive neuron count, five 500μm x 500μm fields per section were randomly chosen in the cortical layer V and analyzed. The average value of 5 fields was presented for each mouse.

### Bulk RNA-Seq

Fifteen thousand cells were sorted out into TRIzol and stored at-70°C until library preparation. Total RNA was purified by phase extraction and processed to cDNA using SMART-Seq v4 Ultra Low Input RNA Kit (Takara bio USA, CA) with 15-cycle amplification. Adaptors were added to 1ng cDNA using the Nextera XT library prep kit (Illumina, CA). The libraries were sequenced on Nextseq500 with 55bp paired end read. For sequence data analysis, adapter sequences were trimmed with Trimmomatic v0.33 after the initial quality check, and then data was aligned with STAR v2.5.2b. Sequence duplicates were removed using Picard v2.17.11. Aligned tags were counted with Subread featureCounts, and differentially expressed genes were estimated using DESeq2^51^ with FDR<0.01. Batch effects were normalized at this point if needed. Heatmaps were depicted with Morpheus (https://software.broadinstitute.org/morpheus).

### Single-cell RNA-seq for constitutive Irf8 knockout microglia

Ten thousand cells were sorted out from 3 mice pools and immediately emulsified and subjected to the libraries using a 10x Genomics Chromium kit v2 as the manufacturer’s protocol. The libraries were sequenced 26bp for R1 and 98bp for R2 on HiSeq2000. The fastq files were first processed with Cellranger v3.1 pipeline to make gene expression matrices, and the following analyses were performed with Scanpy^52^. Cells with less than 500 or greater than 6,000 detected genes were filtered from analysis. Genes detected in less than three cells were also removed from the data. The feature counts were normalized to 10,000 reads per cell and logarithmized. The data were regressed out based on the highly variable genes marked with *min_mean=0.0125, max_mean=3, min_disp=0.5* options, and the cells exhibiting high standard deviation (SD>15) were excluded from the analysis. UMAP dimension reduction and Leiden clustering were employed to visualize the data with *n_neighbors=80, n_pcs=15,* and *min_dist=1.0* parameters. Each cluster’s connectivity was computed using PAGA^42^ with the default parameters.

### Single-cell multiome for conditional Irf8 knockout microglia

Forty thousand male microglia were isolated from each mouse and immediately emulsified and subjected to the libraries using a Chromium Next GEM Single Cell Multiome ATAC + Gene Expression Reagent as the manufacturer’s protocol. The ATAC libraries were sequenced 50bp for R1, 8bp for the i7 index, 24bp for the i5 index, and 49bp for R2 on NovaSeq6000. The RNA libraries were sequenced 28bp for R1, 10bp for i7, 10bp for i5, and 90bp for R2. The fastq files were processed with Cellranger-arc pipeline to create a single sequence aggregate, and the following analyses were performed with Muon^53^.

Cells with low-quality sequences were filtered from the analysis. To that end, the following criteria were used: less than 200 detected transcripts, less than 500 unique molecular identifiers (UMIs), or >1.0% mitochondrial gene contamination for scRNA-seq, peaks within less than 10 cells, cells with less than 1000 or greater than 80,000 UMIs, cells with less than 100 or greater than 30,000 peak regions detected, for scATAC-seq. The feature counts were log-normalised, the same as the above scRNA-seq analysis. The data were regressed out based on the highly variable genes marked with *min_mean=0.02, max_mean=4, and min_disp=0.5* options. ATAC highly variable counts were filtered with the Cellranger flavor option. For 14,391 cells, UMAP dimension reduction and Leiden clustering were employed to visualize the data with *n_neighbors=100, n_pcs=50,* and *min_dist=1.0* parameters for scRNA-seq, *n_neighbors=100, n_pcs=30,* and *min_dist=1.0* for scATAC-seq. The transcriptome and chromatin accessibility data with identical barcodes were then integrated. The differentially accessible peaks were identified by the student’s t-test combined with Benjamini-Hochberg correction, and peaks with adjusted p-value < 0.01 were considered statistically significant. HOMER was used for motif analysis and peak overlap identification. We utilized ArchR^54^ to generate pseudobulk bigwigs, which were normalized based on the number of fragments.

### CUT&RUN

CUT&RUN methodology^55^ was used with some modifications. Protein A-bound micrococcal nuclease (pA-MNase) was kindly provided by Steven Henikoff. For histone marks, live microglial cells were sorted out and washed once with HEPES-NaCl buffer (HEN buffer; 20mM HEPES pH7.5, 150mM NaCl, 500µM Spermidine, Protease inhibitors cocktail), then counted. Ten thousand cells were immobilized to activated BioMag®Plus Concanavalin A (ConA)-beads (Bangs Laboratories, IN) in the presence of Mn^2+^ and Ca^2+^ ions (0.1mM MnCl_2_, 0.1mM CaCl_2_) at room temperature. The beads complex was then permeabilized using 0.005% Digitonin (Merck)-containing HEN buffer (Digitonin buffer) and incubated for 12hrs at 4°C with 0.25µg of the following antibodies in the presence of 2mM EDTA: rabbit anti-H3K27ac (ab4729, Abcam), rabbit anti-H3K4me1 (ab8895, Abcam). After antibody binding, beads were washed twice with Digitonin buffer and bound pA-MNase. After the washout of extra pA-MNase, beads-immunocomplex was activated in the presence of 3mM CaCl_2_ at 0°C for 30 min. The digestion was stopped by adding 10x Stop buffer (1.70M NaCl, 100mM EDTA, 20mM EGTA, 0.25mg/ml Glycogen, 0,25µg/µl RNaseA) and 15pg MNase-digested yeast DNA spike-in control. The nuclei were digested at 56°C for five hours after adding 40µg of Proteinase K and SDS at the final concentration of 0.1%. The immunoprecipitated DNA was purified by phase separation and removed undigested DNA with a half volume of AMPure beads. For library preparation, purified DNA was processed using SMARTer® ThruPLEX DNA-Seq Kit (TAKARA Bio USA, CA) as the manufacturer’s protocol except for PCR cycles to avoid amplifying long fragments (13 cycles, 10s for extension). For transcription factors, the antibody-labeled brain homogenate was fixed with 1% Paraformaldehyde-PBS at room temperature for 10 minutes right before cell sorting. Sorted cells were immediately lysed with nuclear extraction buffer (20mM HEPES pH7.5, 10mM KCl, 500µM Spermidine, 0.1% Triton X-100, 20% Glycerol, Protease Inhibitor cocktail) and counted. Thirty-thousand nuclei were immobilized with ConA-beads in the same nuclear extraction buffer and blocked using Wash buffer (20mM HEPES pH7.5, 150mM NaCl, 500µM Spermidine, 0.1% Bovine Serum Albumin, 1% Triton X-100, 0.05% SDS, Protease inhibitor cocktail) containing 4mM EDTA. The nuclei were incubated with a rabbit anti-GFP antibody (0.5µg/reaction, ab290, Abcam) or a rabbit anti-mouse PU.1 (EPR22624-20, Abcam) for 12hrs at 4°C. After binding to pA-MNase in Wash buffer, DNA was cleaved by adding 3mM CaCl_2_ for 30min at 0°C. MNase was inactivated by adding chelators mixture at the final concentration of 170mM NaCl, 10mM EDTA, 2mM EGTA, 0.005% NP-40, 25µg/ml Glycogen, and 15pg yeast spike-in control. Total DNA was extracted, and de-crosslinked by incubating overnight at 65°C in the high-salt buffer containing 0.1% SDS, 0.5mg/ml Proteinase K and 0.1mg/mL RNase A. The undigested DNA was removed as above, and the small fragments for sequencing libraries were harvested. The final libraries were purified by AMPure beads and sequenced with a 30bp paired end read. For data analysis, the adapter and low-quality sequences were removed from raw fastq data using TrimGalore. Then the data was aligned on mm10 mouse genome using Bowtie2 with *--local--very-sensitive-local--no-unal--no-mixed--no-discordant--phred33-I 10-X 700--no-overlap--no-dovetail* options. Peak calling was performed by MACS2 with *--broad--keep-dup all-p 0.01--min-length 1000* option for a broad histone mark, *--keep-dup all-q 0.01 –max-gap 640* option for a narrow histone mark and IRF8^56^. For PU.1, we employed *-q 0.05* instead because it is more consistent with the previous dataset (GSM1533906)^33^. The peak co-occurrence, genomic position, and motif analysis were performed using HOMER^34^ with *-d given* option. To determine highly concordant peaks across biological replicates, we employed “the Majority rule,” which picks up only peaks found within more than 50% of replicates^57^. The differential peaks were obtained using Diffbind-DESeq2^58^ with FDR<0.05 cut-off. Then all bam files were merged and used for IGV visualization and depicting average plots and heatmaps with Deeptools^59^. H3K27ac^high^ large regions were identified as “super-enhancer regions” of the ROSE’s readouts with default parameters^35^.

### Bulk ATAC-Seq

ATAC-Seq libraries were constructed as described before^37^. Three thousand (X04 microglia) to fifty thousand isolated microglia were lysed in lysis buffer (10mM Tris-HCl pH7.4, 10mM NaCl, 3mM MgCl_2_, 0.1% NP-40). Following lysis, nuclei were tagmented using the Nextera DNA library prep kit (Illumina) for 30 minutes. Then, DNA was purified and amplified for 9-14 cycles based on the SYBR-qPCR determination. The libraries were sequenced with a 50bp paired end read. After removing the adapter and low-quality sequences, the data were aligned to the mm10 genome with Bowtie2. Peaks were called on aligned sequences using MACS2 with *-p 0.01--nomodel--shift-37--extsize 73-B--SPMR--keep-dup* all options. Peaks were then evaluated with less than 0.01 irreversible differential rates. Differentially accessible regions were identified using Diffbind-EdgeR with FDR<0.05^58^. Deeptools and HOMER were employed for further analyses, including motif enrichment analysis with the indicated options.

### Whole-genome bisulfite sequence

Genomic DNA was extracted from 40,000 male microglial cells using Quick-DNA Microprep plus kit (Zymo Research) as the manufacturer’s protocol. Then, 50ng DNA was bisulfite-converted and processed to the library using Pico Methyl-Seq Library Prep Kit (Zymo Research), with 8-cycle PCR amplification. The libraries were sequenced with 75bp paired end read on HiSeq3000 in the NHLBI sequencing core (9.2x coverage on average). Based on the Bismark M-bias plot, the adapters, low-quality sequence, and an additional 10bp of 5’-end and 2bp of 3’-end were removed from the sequencing data using TrimGalore. The data were first aligned using Bismark^40^ with *-q--score-min L,0,-0.4--ignore-quals--no-mixed--no-discordant--dovetail--maxins* options, and then unmapped sequences were aligned additionally with single-end mode. After deduplication and methylation calls, both files were merged into a single coverage file. Differential analyses were performed using R/DSS package^41^ using FDR<0.05 for the differentially methylated loci (DMLs) and p<0.001 for the regions (DMRs).

## Data availability

All high-throughput sequence datasets generated in this paper are available in GSE231406 and GSE266424. Some gene sets were obtained from publications as denoted individually.

## Abbreviations

AD: Alzheimer’s disease
DEG: differentially expressed gene
FACS: fluorescence-activated cell sorting
FC: fold change
FDR: false discovery rate
GFP: green fluorescent protein
GO: gene ontology
IRF8: interferon regulatory factor 8
TF: transcription factor
TES: transcription end sites
TSS: transcription start sites
YFP: yellow fluorescent protein
WT: wild type

## Acknowledgments

We thank Steven Coon and James Iben, NICHD DIR MGC, for their guidance for scRNAseq and multiome, Yuesheng Li in NHLBI sequencing core for whole-genome bisulfite sequencing, and Fayaz Seifuddin for data analysis. We also thank Avinash Bhandola for providing Rosa26-EYFP mice, Lisa William-Simons for cell sorting, and the NICHD animal facility staff for caring for our mice and for other technical assistance. We gratefully acknowledge colleagues in the NIH and elsewhere for valuable advice and discussions. This work was supported by the NICHD Intramural programs ZIA HD008015-13.

## Author Contributions

K.S and K.O conceived the framework of this work; K.S generated IRF8cKO microglia, devised whole genome analyses, and executed experiments along with R.P; E.L performed immunohistology; D.K. provided advice on experiments; K.S and K.O wrote the manuscript.

