## Supplemental Figures and Text for "IRF8 configures enhancer landscape in postnatal microglia and directs microglia specific transcriptional programs"

Supplemental Figure Legend

Supplemental Figure 1

A. Gating strategy for the isolation of microglia and peritoneal macrophages. Cells were gated with $CD11b^{+}CD45^{low}$ for microglia and $CD11b^{+}F4/80^{+}$ for peritoneal macrophages. We used backgating to enrich the target cells efficiently. All FACS profiles shown were after backgating. Staining with 7-AAD was performed when the live cells were needed.
B. IRF8 expression in microglia and peritoneal macrophages. Flow cytometry was performed to detect GFP for cells from IRF8-GFP/+ hemizygous transgenic mice (filled in magenta) and WT control (black line). The GFP expression in P9 microglia and P14 microglia of IRF8-GFP/GFP mice are also shown on the right.

Supplemental Figure 2

A. Flow cytometry of 7-AAD-negative brain cells stained with CD11b and CD45 antibodies from WT and IRF8KO mice. The median ratio of each subset from 7-AAD-negative cells was shown.
B. Absolute cell number of brain myeloid cells. The number of $CD11b^{+}CD45^{low}$ cells for microglia, $CD11b^{+}CD45^{low}Ly6C^{+}$ for P3 cells, $CD11b^{+}CD45^{low}Ly6C^{+}CCR2^{+}$ for Monocyte, $CD11b^{+}CD45^{low}Ly6G^{+}$ for Granulocytes were presented.
C. Flow cytometry profiles of CX3CR1, Ly6C, F4/80, and TMEM119 expression (filled in magenta) and isotype IgG (black line) in microglia from WT and IRF8KO brains or P3 population from IRF8KO brains.

D. Flow cytometry of 7-AAD$^-$CD11b$^+$CD45$^+$ brain cells stained with CD206 and Ly6C antibodies from WT and IRF8KO mice. P3 cells (represented by Ly6C$^+$ cells in the lower right quadrant) in the IRF8KO brain do not express the BAM cell marker CD206.

E. Flow cytometry of 7-AAD$^-$CD11b$^+$CD45$^+$ brain cells from P7 and adult stained with two BAM cell markers, CD206 and FOLR2 antibodies, WT and IRF8KO mice to analyze BAM cells. The BAM cells occupied less than 1.0% of total cells in any case.

F. The absolute cell number of BAM cells calculated using FACS data in Fig.S2E.

G. Microglia identity gene expression in P9, P14, and adult WT microglia

Supplemental Figure 3 (also see Method)

A. MA plots showing H3K27ac (left) and H3K4me1 (right) histone marks comparing WT and IRF8KO microglia (n=2). The differential regions with FDR<0.05 are colored in pink. The regions found only in WT microglia were labeled WT-specific, while KO-specific stands for the regions absent in WT.

B. MA plots showing PU.1 peak intensity comparing WT and IRF8KO microglia (n=2). The differential regions with FDR<0.05 are colored in pink.

C. The distance between two neighboring IRF8 peaks within an H3K27ac high region: Median;11.418 kb, 75% Quantile; 38.977 kb.

D. Venn diagram showing the overlap between H3K27ac$^{high}$ regions and regions occupied with an array of enhancer IRF8 peaks. To define an array of enhancer IRF8 peak regions, H3K4me1 marked IRF8 peaks within 40kb intervals were concatenated by ROSE. IRF8 also forms an array of bindings outside $H3K27ac^{high}$ regions (IRF8nSEC). See more details in the Supplemental methods below.

E. Genomic size of $H3K27ac^{high}$ regions (super enhancers) and IRF8nSEC regions (smaller enhancers).

F. Genome wide distribution of super enhancers and IRF8nSEC region determined with HOMER.

G. Representative IGVs. Orange bar; IRF8nSEC region (smaller enhancers).

Supplemental Figure 4

A. M-A plot showing the differential ATAC signals comparing P9 and P14 microglia (left; n=2-3). The differential regions with FDR<0.05 are colored in pink. The gained regions occupied most of the newly bound IRF8 at P14 (right Venn diagram). De novo motif analysis for the overlapping area in the Venn diagram (bottom) indicated the potential transcription factors that co-bind to newly bound IRF8.

B. M-A plot showing the differential ATAC signals comparing P14 and Adult microglia (left; n=2). The differential regions with FDR<0.05 are colored in pink. The gained regions were considerably occupied with newly bound IRF8 in adults (right Venn diagram). De novo motif analysis for the overlapping area in the Venn diagram (bottom) indicated Batf3 and PU.1 may co-bind to newly bound IRF8 in adults.

C. Correlation biplot showing ATAC signal intensity and DNA methylation levels in 1,337 regions where all differential ATAC peaks and DMRs overlapped. Analyzed with the Kendall rank correlation test (coefficient r=-0.75, $p=3.28x10^{-229}$).

D. Gene ontology for DMRs with GREAT. The top 5 terms with corresponding binomial FDR values were shown.

Supplemental Figure 5

A. Overlap of DEGs between bulk RNA-Seq and scRNA-Seq used in this paper.

B. GO analysis of 548 genes enriched in IRF8KO Cluster 4 compared to the other IRF8KO clusters. The top 5 categories are shown. See Supplemental Methods for identifying DEGs from scRNA-seq data.

C. UMAP depicting a population expressing Ly6c2 mRNA

Supplemental Figure 6

A. Gel electrophoresis showing Cre-excision PCR products using genomic DNA of sorted WT ($Irf8^{flox/flox}$) or IRF8cKO ($Irf8^{flox/flox}Cx3cr1^{CreERT2}$) microglia. The data from two different Tamixifen starting dates (left; P12, right; P28) were presented. In each regimen, Tamoxifen was injected for 5 consecutive days, and YFP-expressing microglia were analyzed. The $\mathit{Cx3cr1}^{\mathit{CreERT2}}$ genotyping for the same cells was shown on the right. The band indicated "not excised" was also detected in IRF8cKO microglia. M; 100bp DNA ladder

B. Volcano plot for RNA-seq comparing WT and IRF8cKO microglia. Genes upregulated in IRF8cKO microglia over WT are colored pink (n=546), or genes downregulated in WT are colored blue (n=334, FDR<0.01).

C. Overlap of DEGs between bulk IRF8KO RNA-seq and IRF8cKO RNA-seq data.

D. Gene set expression in bulk WT and IRF8cKO microglia (compare with heatmaps in Fig. 2B). A part of microglia identity genes and CLEAR lysosomal genes were downregulated in IRF8cKO microglia, while some interferon-related and DAM-like genes were expressed higher in IRF8cKO microglia.

Supplemental Figure 7

A. Volcano plot for RNA-Seq comparing WT and 5xFAD/WT microglia. Genes upregulated in 5xFAD/WT microglia over WT are set as a 5xFAD$^{-}$associated gene set, colored in pink (n=699, FDR<0.08). or genes downregulated in WT are colored in blue (n=137). Representative genes are labeled.

B. Flow cytometry of 9-months-old, live CD11b$^{+}$CD45$^{+}$ brain cells stained with MethoxyX04 (MetX04) and Tmem119 antibodies from WT (no 5xFAD control) and 5xFAD/WT mice for bulk ATAC-seq. The ratio of X04$^{+}$ and X04$^{-}$ subsets was shown. We isolated WT X04$^{-}$, 5xFAD/WT X04$^{-}$, and 5xFAD/WT X04$^{+}$ cells to perform ATAC-seq (n=2).

C. Flow cytometry of 6-month-old male CD11b$^{+}$CD45$^{+}$ brain cells stained with MetX04 and Tmem119 antibodies from 5xFAD/WT and 5xFAD/IRF8KO mice. The ratio of X04$^{+}$ microglia in each genotype was quantified on the right (n=4-5). Data: Mean +/- SEM; *: p-value <0.05 by unpaired student's t-test.

D. Heatmaps presenting scATAC-seq signal intensity in the DAM gene regions of WT and IRF8cKO microglia clusters (left), and that of bulk ATAC-seq in 5x FAD X04$^{+}$, X04$^{-}$, WT, and IRF8KO microglia (right). To evaluate the correlation between X04$^{+}$ microglia and IRF8cKO with respect to DAM genes, we set the regions identified in Fig. 6J which were linked to DAM gene expression in IRF8cKO microglia. In sc ATAC-seq, IRF8cKO microglia clusters 2 and 3 exhibited a higher signal level than WT microglia clusters. In bulk, IRF8KO microglia showed a substantial increase of ATAC-seq signals in those regions compared to WT microglia, whereas X04$^{+}$ microglia showed no increase of the signal compared to X04$^{-}$ microglia, suggesting a distinct chromatin profile in X04$^{+}$ microglia compared to that in IRF8cKO microglia.

E. Representative histology images depicting immunostaining of the cortex regions of 6-month-old 5xFAD/WT or 5xFAD/IRF8cKO brain with 6E10 for Aβ (white), Iba1 for microglia (red), and Hoechst33342 for nuclei (blue) identification. Merge images on the right illustrate a lack of extensive colocalization in the 5xFAD/IRF8cKO brain. Data: Mean +/- SEM; *: p-value <0.05.

F. Representative histology showing core-dense plaques in the cortex region of four-month-old and one-year-old 5xFAD/WT or 5xFAD/IRF8KO brain. Brain sections were stained with Thioflavin S (green). The number of ThioS$^{+}$ aggregates (right), amount of Aβ deposition (bottom), and size of Aβ plaques (upper left) were quantified and presented alongside. Values represent the average of five fields from each brain and statistically tested by one-way ANOVA with Tukey's post hoc test ($F (3, 33) = 16.17$, $P<0.001$ for the number of aggregates, F (3, 21) = 23.48, P<0.001 for plaque size) . Scale bar: 200 μm. Data: Mean +/- SEM; *: p-value <0.05.

Supplemental Table S1

The list of DEGs in Batf3KO and Sall1KO microglia that overlap with those in IRF8KO microglia, related to Fig.2E and Fig.2F.

Supplemental Table S2

The list of overlapping peaks among IRF8, Sall1, and PU.1 ChIP data, related to Fig. 2G. The nearest gene from each peak identified by HOMER was also listed.

Supplemental Table S3

Potential transcription factors that may co-bind to IRF8 in microglia before P14 and after P14. The HOMER known motif analyses were performed for the overlapping area of the Venn diagram in Fig.S4A and Fig.S4B and presented.

Supplemental Methods

*Identification of non-super enhancer IRF8 cluster regions (IRF8nSEC)*

To find the genomic regions occupied by dense, multiple IRF8 peaks similar to $H3K27^{high}$ regions, we analyzed the intervals of IRF8 peaks within $H3K27^{high}$ regions as shown in Figure S3C. Next, by taking advantage of the 75% Quantile value in Figure S3C, the peaks within 40kb intervals were grouped so that we could specify the regions occupied by IRF8 of similar linear density. To avoid the contamination of promoter IRF8, we removed the IRF8 peaks without H3K4me1 marks and the ones within 2.5kb from TSS in advance. ROSE was employed for grouping IRF8 peaks with *-s 40,000* option instead of *-s 12,500*. The grouped IRF8 cluster regions, except super-enhancer regions, were defined as "non-super enhancer IRF8 cluster regions" (IRF8nSEC). The density of IRF8 peaks and region size were compared to the super-enhancer regions by the Kolmogorov-Smirnov test (Figure S3C, S3D).

*Identification of differentially expressed genes in scRNA-Seq clusters*

The differential expression analysis for each cluster was performed by SigEMD[1], where the number of permutations was set to 100, and genes were filtered with p.adjust<0.01. The consistency between this DEG estimation and bulk RNA-seq transcriptome was examined in Supplemental Figure 5B. The genes enriched in IRF8 Cluster 4 were identified in comparison with the other IRF8 clusters.

*Quantitative PCR to analyze the deletion efficiency by Cre recombinase*

The genomic DNA from five to twenty thousand EYFP-positive microglia was isolated with Quick-DNA Microprep plus kit (Zymo Research) as the manufacturer's protocol. The DNA for the standard curve was prepared similarly from mouse cells. Quantitative PCR was performed using Fast SYBR Green Master Mix (Applied Biosystems) with the following primer set.

Control chr8 region Forward: 5'-TTTCAAGCCTTCTGTCTGCG-3'

Control chr8 region Reverse: 5'-CACCGCGTGCTCAACTACTC-3'

Irf8 exon 2 Forward: 5'-ATCGAACAGATCGACAGCAGC-3'

Irf8 exon 2 Reverse: 5'-CTTCCAGGGGATACGGAACA-3'

*Deconvolution analysis*

MuSiC deconvolution[2] was performed to determine how many BAM cells were contaminated in our bulk RNA-seq datasets, referring to the deposited dataset, GSM6705227. The BAM proportion in each microglia dataset was calculated as below.

$$BAM\ contamination\ (\%) = \frac{\sum(Est.Prop.of\ all\ BAM\ culsters) \times 100}{\sum(Est.Prop.of\ all\ microglia\ clusters)}$$

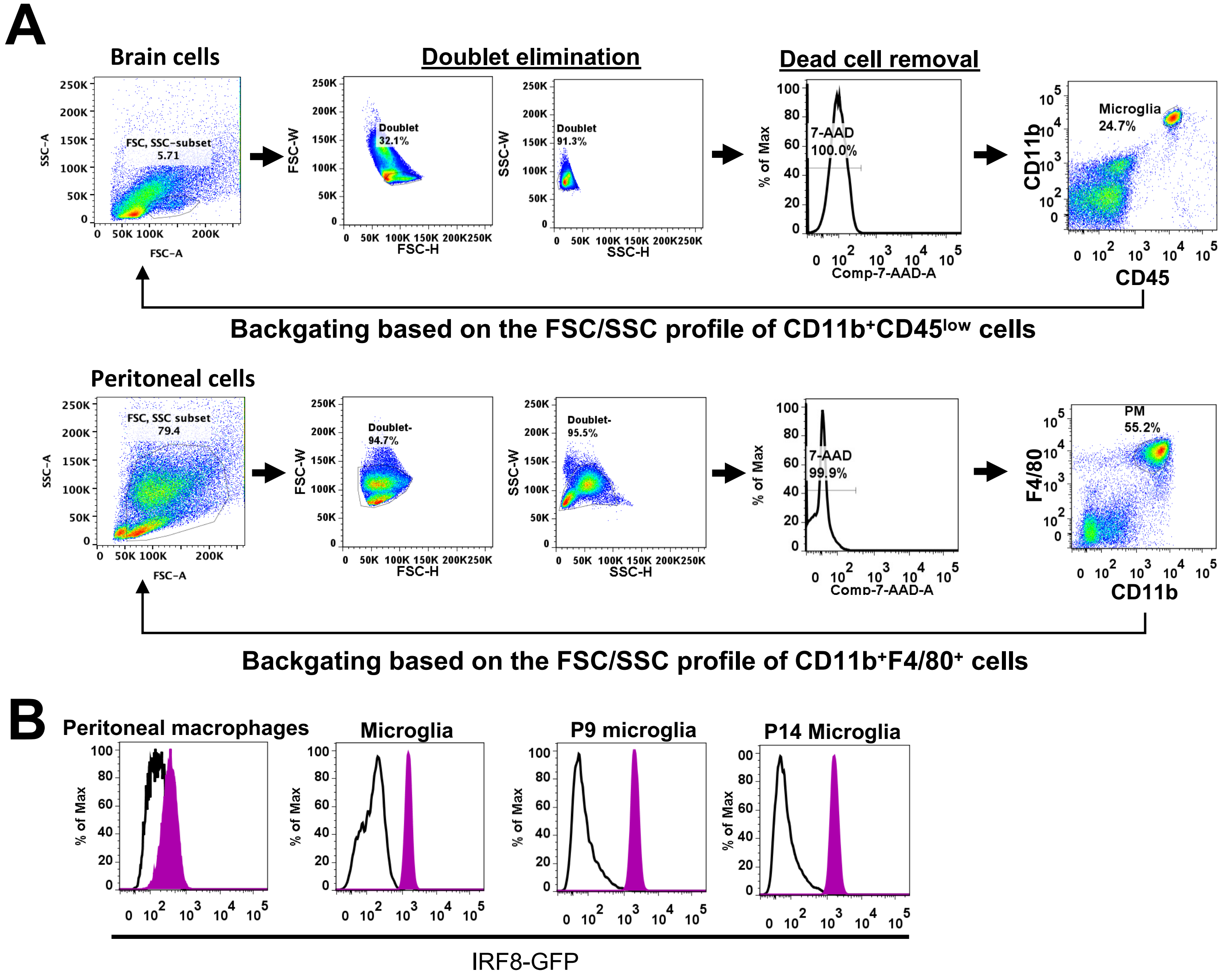


Fig.S1 Saeki et al.

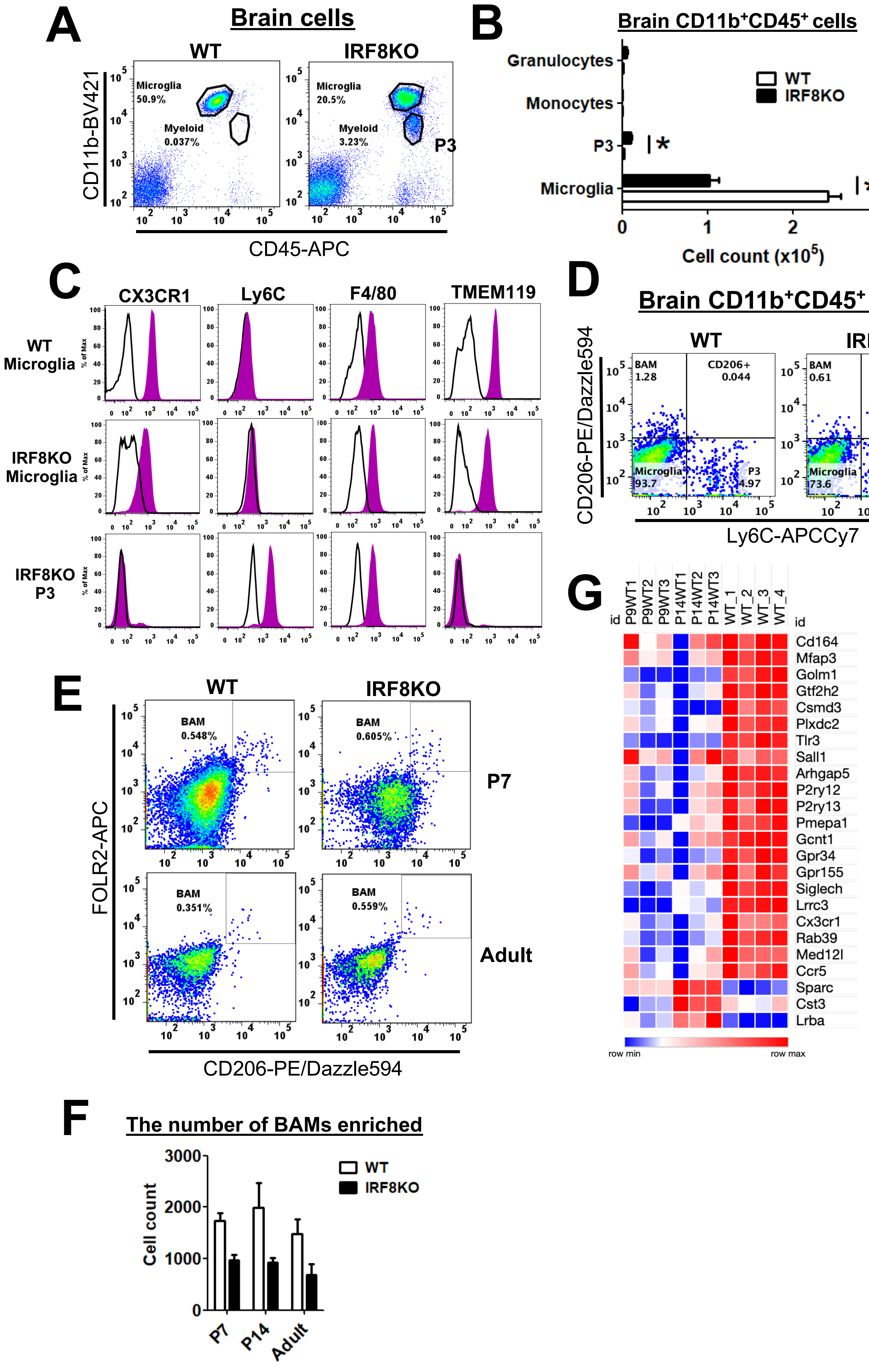


**Fig.S2 Saeki et al.**

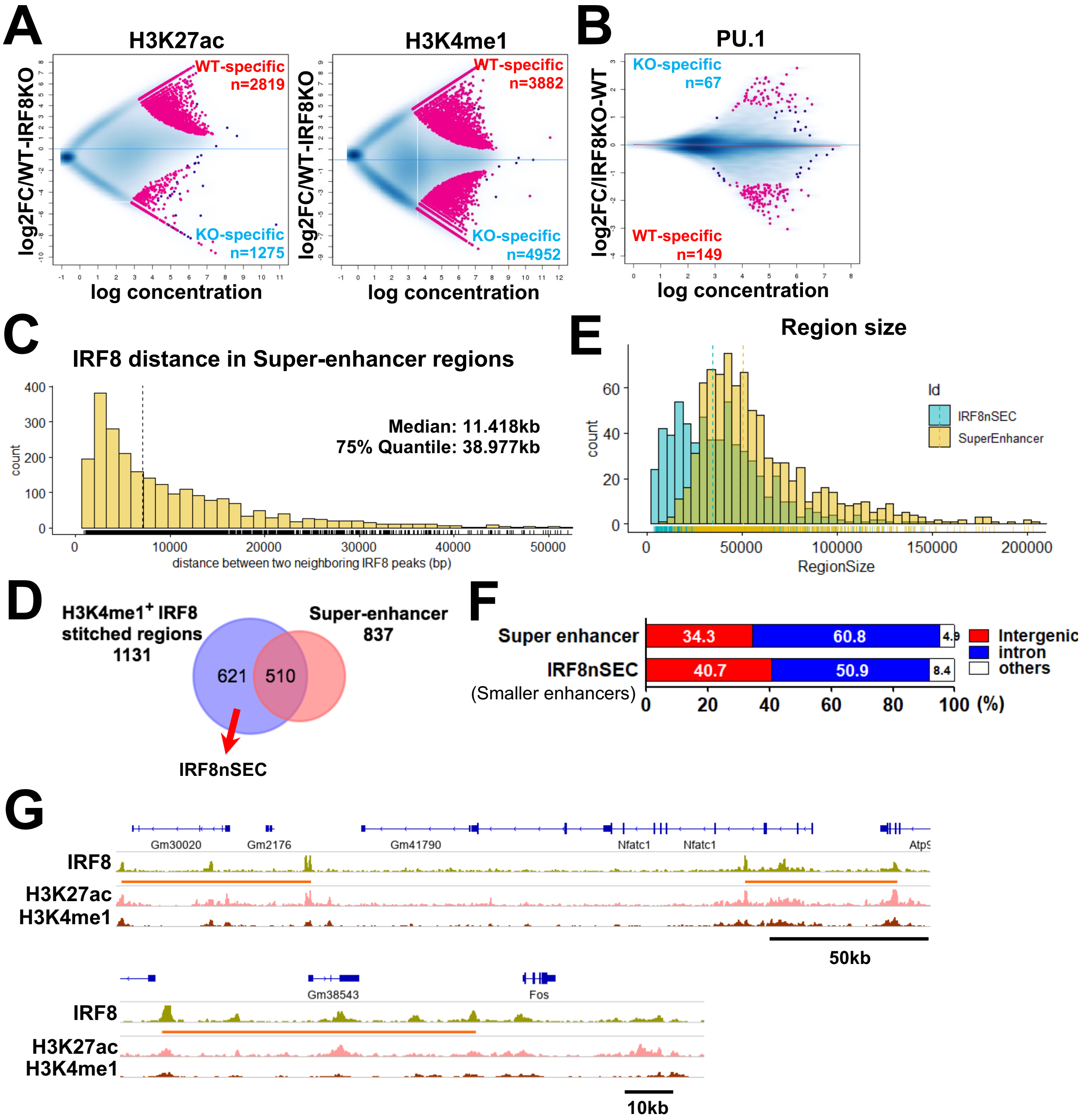


### Fig.S3 Saeki et al.

**A**

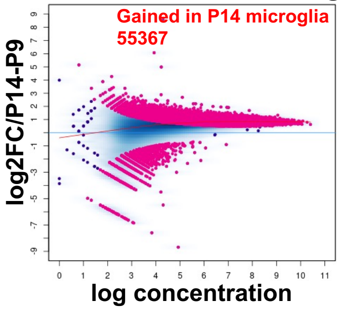


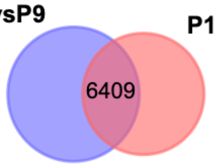


**potential *cis*-regulatory factors co-bound to IRF8 in P14 microglia**

| Rank | Motif | P-value | T/B (%) | TF |
|---|---|---|---|---|
| 1 | 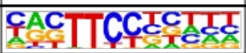 | 1e-1238 | 64.69/20.97 | Sfpi1 (0.921) |
| 2 | 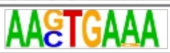 | 1e-286 | 35.89/17.01 | PU.1-IRF8 (0.872) |
| 3 | 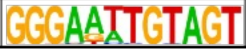 | 1e-40 | 3.11/1.02 | GFY (0.987) |

**B**

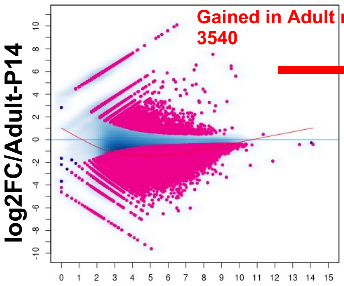


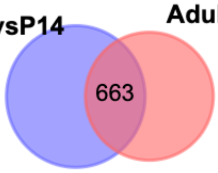


Nearest gene - Ctsc, Tgfbr2 (Batf3 dependent genes)

**potential *cis*-regulatory factors co-bound to IRF8 in Adult microglia**

| Rank | Motif | P-value | T/B (%) | TF |
|---|---|---|---|---|
| 1 | 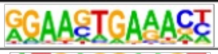 | 1e-95 | 50.68/15.75 | PU.1-IRF8(0.964) |
| 2 | 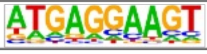 | 1e-36 | 47.96/25.12 | Sfpi1 (0.886) |
| 3 | 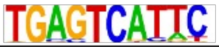 | 1e-13 | 16.89/7.79 | Batf3 (0.904) |

**C**

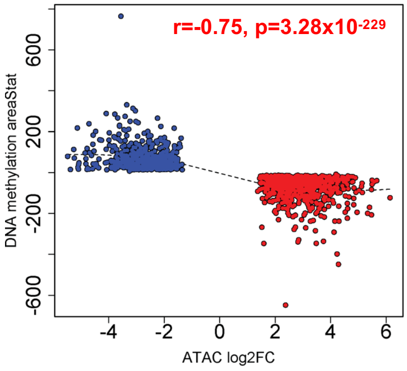


**D**

**Demethylated in WT microglia**

| #Term Name | FDR |
|---|---|
| immune response | 1.26E-15 |
| leukocyte differentiation | 2.49E-13 |
| regulation of cell activation | 2.66E-13 |
| regulation of leukocyte activation | 7.54E-13 |
| adaptive immune response | 7.46E-13 |

**Methylated in WT microglia**

| #Term Name | FDR |
|---|---|
| negative regulation of smoothened signaling pathway | 6.66E-17 |
| positive regulation of Wnt signaling pathway | 1.26E-11 |
| regulation of Wnt signaling pathway | 2.47E-11 |
| regulation of smoothened signaling pathway | 4.76E-11 |
| embryonic digestive tract development | 7.05E-11 |

### Fig.S4 Saeki et al.

A

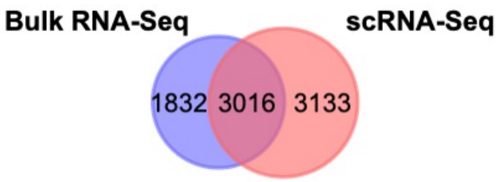


B

**IRF8_CL4 vs rest of IRF8KO microglia**

| Term | Description | FDR |
|---|---|---|
| GO:0009615 | response to virus | -50.564 |
| GO:0035456 | response to interferon-beta | -35.296 |
| mmu05164 | Influenza A - Mus musculus (house mouse) | -12.793 |
| GO:0034341 | response to type II interferon | -11.792 |
| GO:0060759 | regulation of response to cytokine stimulus | -11.513 |

C

**IRF8_CL6 specific**

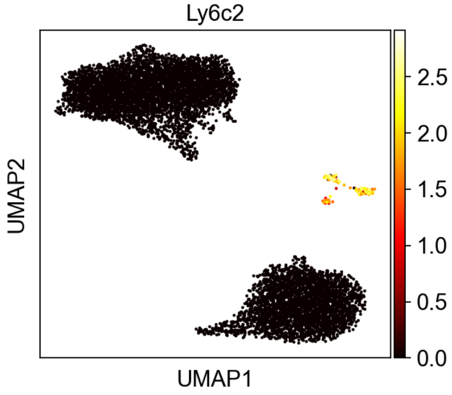


### Fig.S5 Saeki et al.

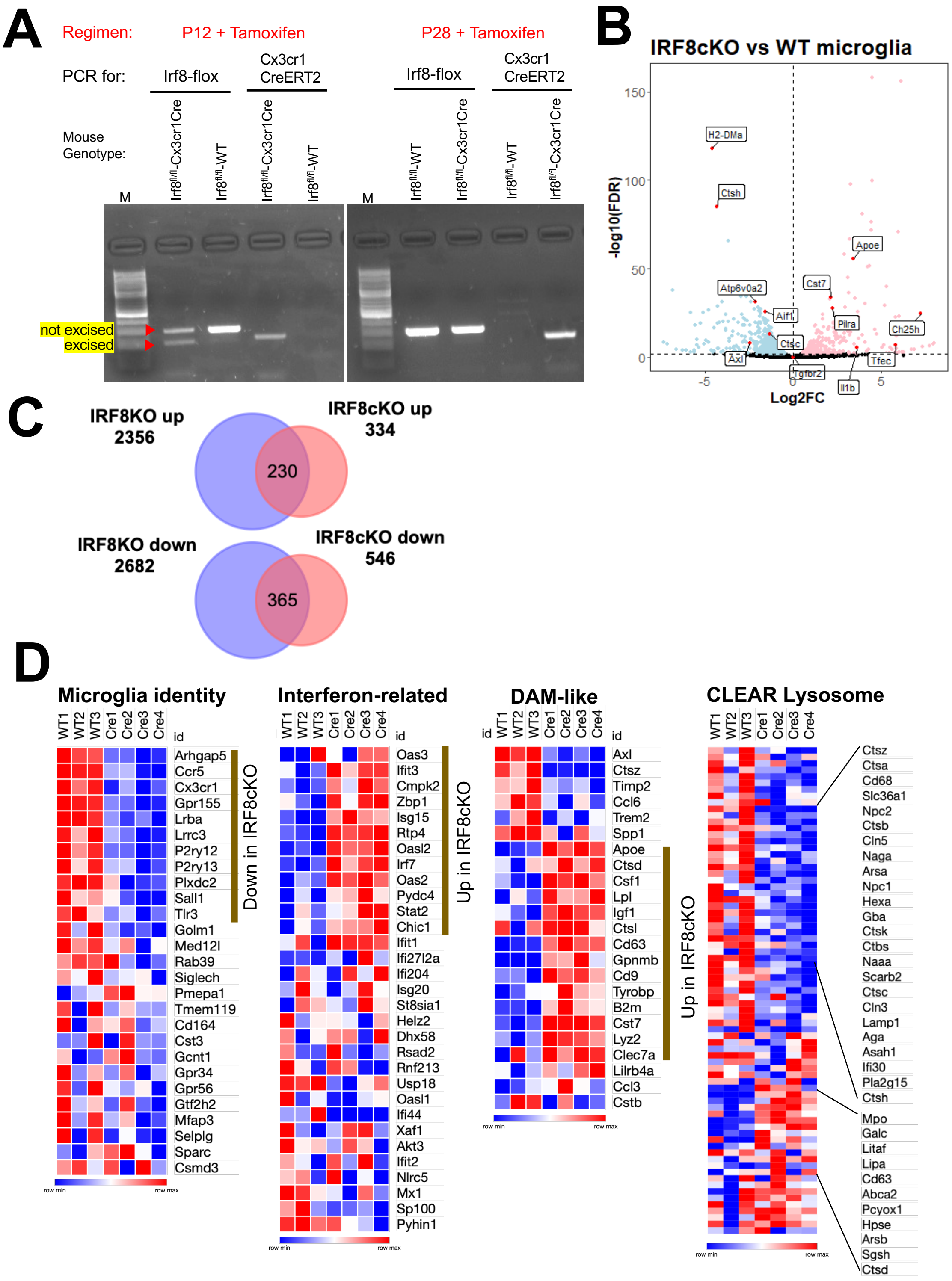


**Fig.S6 Saeki et al.**

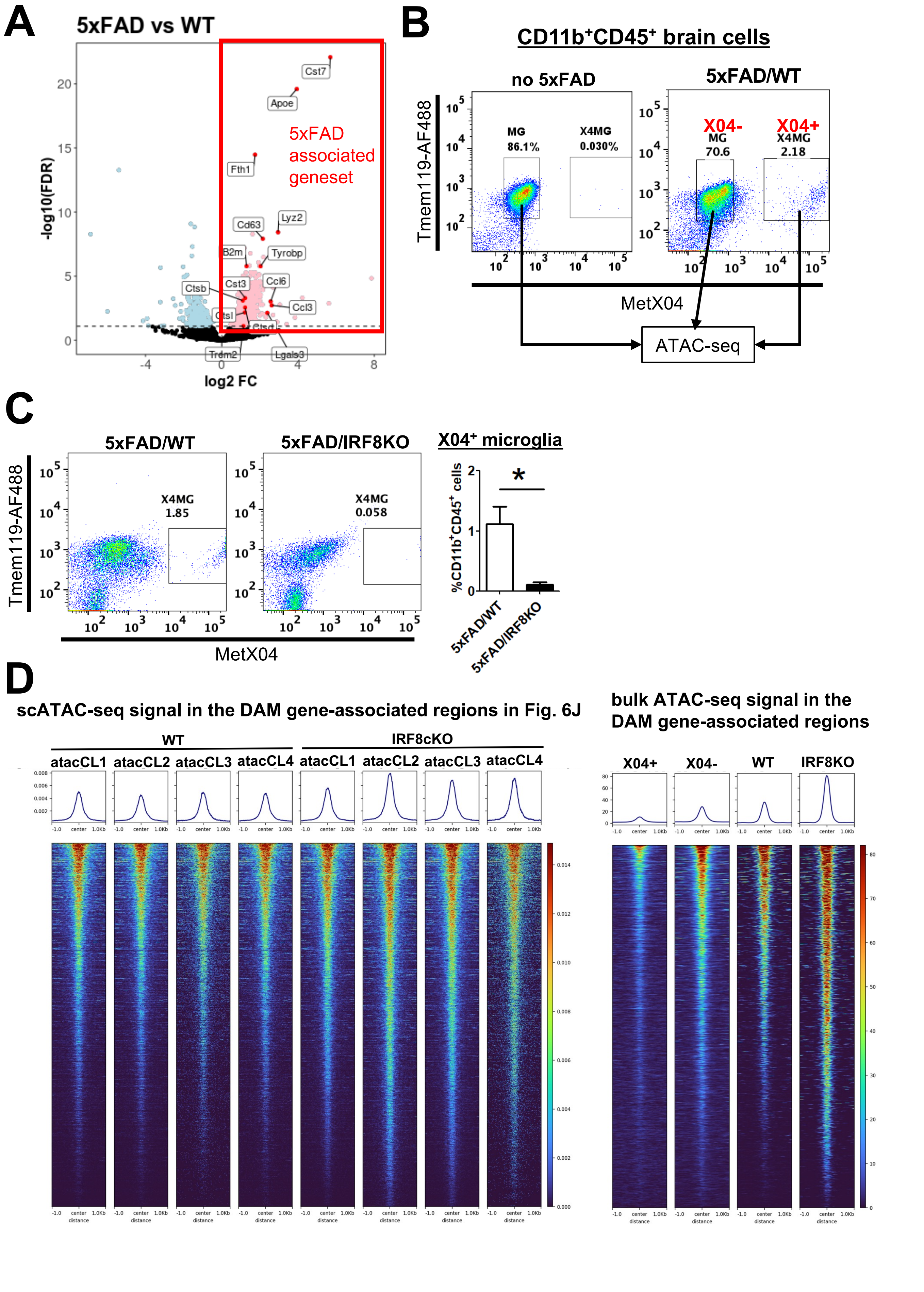


Fig.S7 Saeki et al.

E

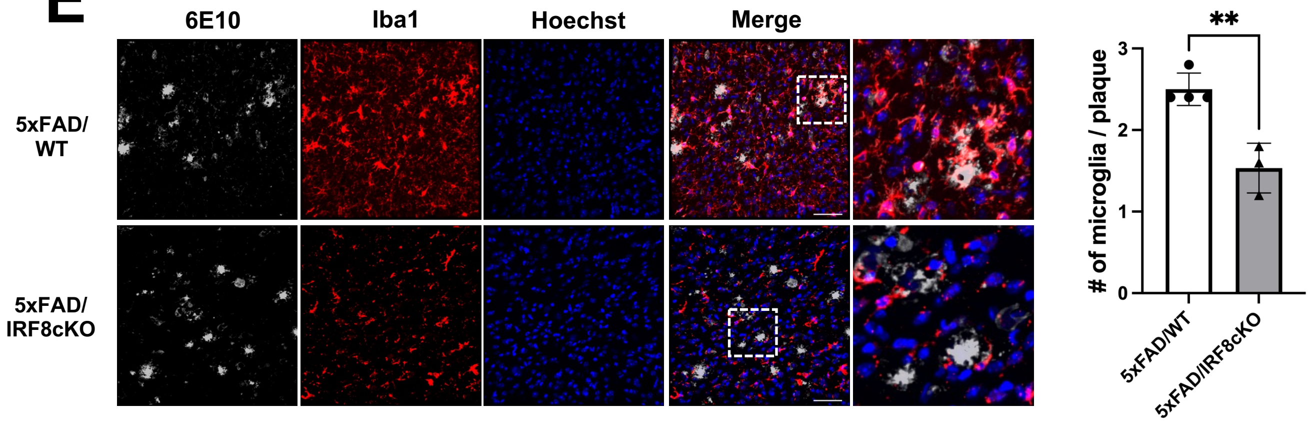


F

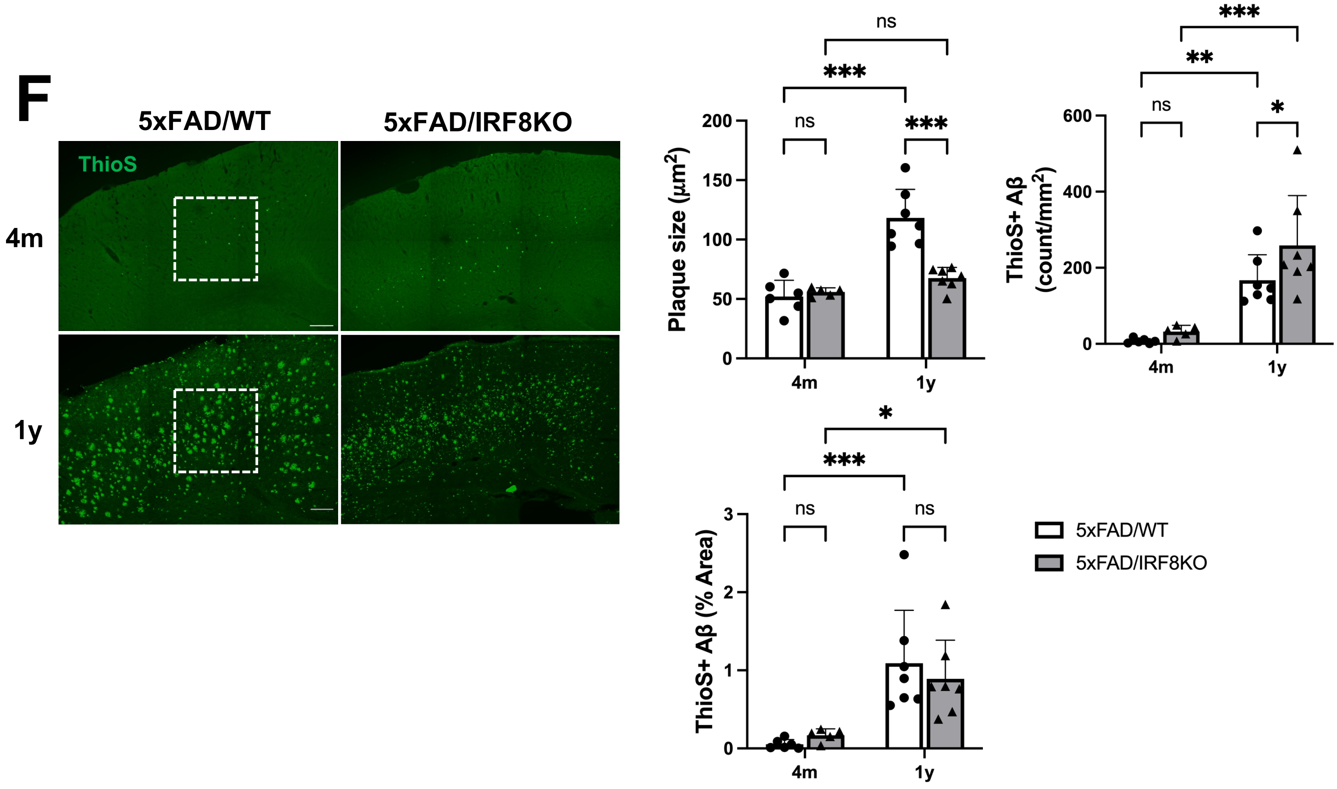


**Fig.S7 Saeki et al.**
